# Convergent behavior of extended stalk regions from staphylococcal surface proteins with widely divergent sequence patterns

**DOI:** 10.1101/2023.01.06.523059

**Authors:** Alexander E. Yarawsky, Andrea L. Ori, Lance R. English, Steven T. Whitten, Andrew B. Herr

## Abstract

*Staphylococcus epidermidis* and *S. aureus* are highly problematic bacteria in hospital settings. This stems, at least in part, from strong abilities to form biofilms on abiotic or biotic surfaces. Biofilms are well-organized multicellular aggregates of bacteria, which, when formed on indwelling medical devices, lead to infections that are difficult to treat. Cell wall-anchored (CWA) proteins are known to be important players in biofilm formation and infection. Many of these proteins have putative stalk-like regions or regions of low complexity near the cell wall-anchoring motif. Recent work demonstrated the strong propensity of the stalk region of the *S. epidermidis* accumulation-associated protein (Aap) to remain highly extended under solution conditions that typically induce compaction or other significant conformational changes. This behavior is consistent with the expected function of a stalk-like region that is covalently attached to the cell wall peptidoglycan and projects the adhesive domains of Aap away from the cell surface. In this study, we evaluate whether the ability to resist compaction is a common theme among stalk regions from various staphylococcal CWA proteins. Circular dichroism spectroscopy was used to examine secondary structure changes as a function of temperature and cosolvents along with sedimentation velocity analytical ultracentrifugation and SAXS to characterize structural characteristics in solution. All stalk regions tested are intrinsically disordered, lacking secondary structure beyond random coil and polyproline type II helix, and they all sample highly extended conformations. Remarkably, the Ser-Asp dipeptide repeat region of SdrC exhibited nearly identical behavior in solution when compared to the Aap Pro/Gly-rich region, despite highly divergent sequence patterns, indicating conservation of function by various distinct staphylococcal CWA protein stalk regions.

## Introduction

The gram-positive bacteria *Staphylococcus aureus* and *S. epidermidis* are of major concern to the healthcare system. Hospital-acquired infections can lead to bacteremia, endocarditis, and prosthetic joint infection. Biofilm formation is a critical virulence factor, which makes these infections difficult to treat [7-10]. It is well understood now that cell wall-anchored proteins, rather than just polysaccharide intercellular adhesin, are extremely important in various stages of biofilm formation and infection [7, 10-12].

Much work has focused on the role of the structurally ordered, functional regions of various cell wall-anchored proteins in biofilm formation [11], however, the regions of low complexity or putative stalk-like regions of these proteins are often neglected. We recently investigated the proline/glycine-rich (stalk-like) region (PGR) of the accumulation-associated protein, Aap, from *S. epidermidis*, and we found this region is intrinsically disordered, but has an unusual ability to remain extended under harsh conditions. Given the placement of this region adjacent to the cell wall, we hypothesize the ability to maintain an extended conformation could help facilitate bacterial surface attachment and intercellular accumulation of staphylococcal cells in a nascent biofilm via the functional adhesive regions of Aap (i.e., the lectin domain and B-repeat superdomain, respectively) [6].

In this study, we sought to understand if such a property was common among similar regions of low complexity from various cell wall-anchored proteins (Figure 1). In addition to the stalk-like PGR region that extends from the bacterial cell wall, Aap features a putatively disordered region at the N-terminus comprised of 11 short A-repeats (Arpts) of ∼16 residues each; this region is included in this study as a control intrinsically disordered polypeptide (IDP) that does not act as a stalk, given that it is not located proximal to the cell wall attachment point [12-14]. The N-terminal ‘A-domain’ of Aap, including the A-repeat region and the downstream lectin domain, is important for surface adherence and adhesion to host cell ligands [15, 16]. The N-terminus must be removed via SepA or other proteases for Aap-dependent biofilm accumulation to occur via Zn^2+^-dependent assembly of the B-repeat superdomain [17, 18]. While the specific role of the A-repeat region is unclear, it may sterically inhibit or modulate intercellular adhesion via the B-repeat superdomain under resting conditions.

**Figure 1.**
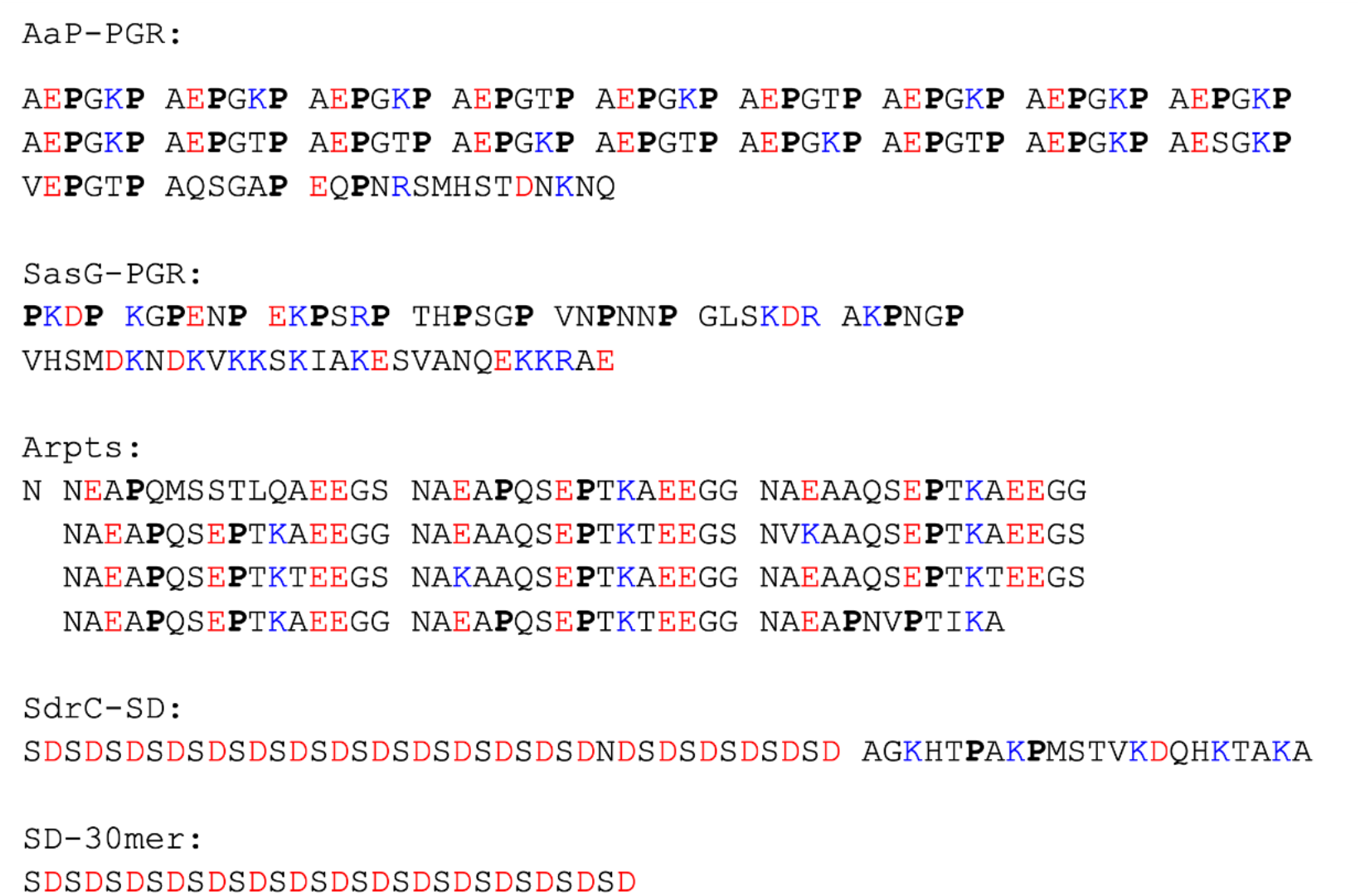
Sequences of IDP constructs. Prolines are in bold font, negatively charged residues are colored red, and positively charged residues are colored blue. Sequence repeats are separated by spaces to highlight the pattern.

Also included in this study is the stalk region equivalent to the PGR of Aap found in the *S. aureus* ortholog, SasG [19]. We hypothesize that SasG-PGR will share very similar properties as Aap-PGR, as both Aap and SasG PGRs contain a proline in every third position (Figure 1) and have similar roles in biofilm formation. In both proteins, a highly extended stalk would likely facilitate the biological functions of host cell interaction as well as intercellular adhesion in the nascent biofilm after cleavage of the A-domain.

SdrC belongs to the serine-aspartate family of proteins expressed by *S. epidermidis* and *S. aureus* [11, 20]. Like Aap and SasG, these proteins are cell wall-anchored and play important roles in biofilm formation, including primary attachment to host tissue via receptor binding and bacterial accumulation via homophilic interaction [11, 20]. The common scheme of serine-aspartate (Sdr) proteins is a large A-region which contains multiple IgG-like repeats involved in ligand binding or homophilic interaction, followed by several B-repeats (unrelated to the B-repeats of Aap and SasG) of ∼110 residues that are elongated in structure and are capable of binding ligands such as collagen [21, 22]. Downstream of the B-repeats is the serine-aspartate repeat region, which contains a large but varying number of SD dipeptide repeats, from 56 residues in SdrG to 558 in SdrF [20]. The C-terminus of the precursor protein contains an LPXTG cell wall-anchoring motif and a hydrophobic membrane-spanning region; as with Aap and SasG, the LPXTG motif is cleaved by sortase A between the T and G residues and the threonine is covalently attached to peptidoglycan in the cell wall [20]. We hypothesize that the SD repeats of SdrC and other Sdr family members will show similar features to the PGR of Aap and SasG, in that they may be highly extended in order to traverse the peptidoglycan layers and to allow the A and B regions to extend outward for efficient binding to host tissues or accumulation via homophilic interactions. Interestingly, the SD repeats lack the high proline content we hypothesized to be important for the extended configuration of Aap-PGR to resist compaction, but they may instead rely on charge-based contributions that are lacking in Aap-PGR.

We utilized several sequence-based predictions that indicated each construct is likely to be disordered, primarily due to the lack of hydrophobic (order-promoting) residues, but in the case of the SD repeats, there is also a strong contribution from high net charge. Predictions based on charge patterning and polyproline type II helix (PPII) propensity suggest highly extended conformations may be preferred by these sequences. Indeed, sedimentation velocity analytical ultracentrifugation (AUC) indicated high frictional ratios representative of highly extended or elongated species, which was confirmed with size-exclusion chromatography (SEC) and small-angle X-ray scattering (SAXS). A battery of experiments using circular dichroism (CD) spectroscopy revealed trends in the resistance to compaction which were proportional to predicted PPII propensity (for proline-rich sequences) or to high negative charge density (for SdrC-SD); specifically, these sequences showed stronger resistance to perturbation by temperature or cosolvents.

## Results

### All constructs are predicted to be disordered

Our first approach to compare these constructs is to examine sequence-based algorithms to determine their propensity to be folded (ordered) or unfolded (disordered). Uversky, et al. found that the mean net charge and mean scaled hydropathy of a protein is a powerful predictor of whether or not the protein will be ordered or disordered [3]. In Figure 2A, all constructs of interest are located on the disordered side of the Uversky plot. Interestingly, while the mean scaled hydropathy values all fall within the range of 0.25 - 0.35, the absolute mean net charge varies much more, ranging from 0.05 to 0.50. In light of the variability in mean net charge, we sought to examine additional parameters related to charge.

**Figure 2.**
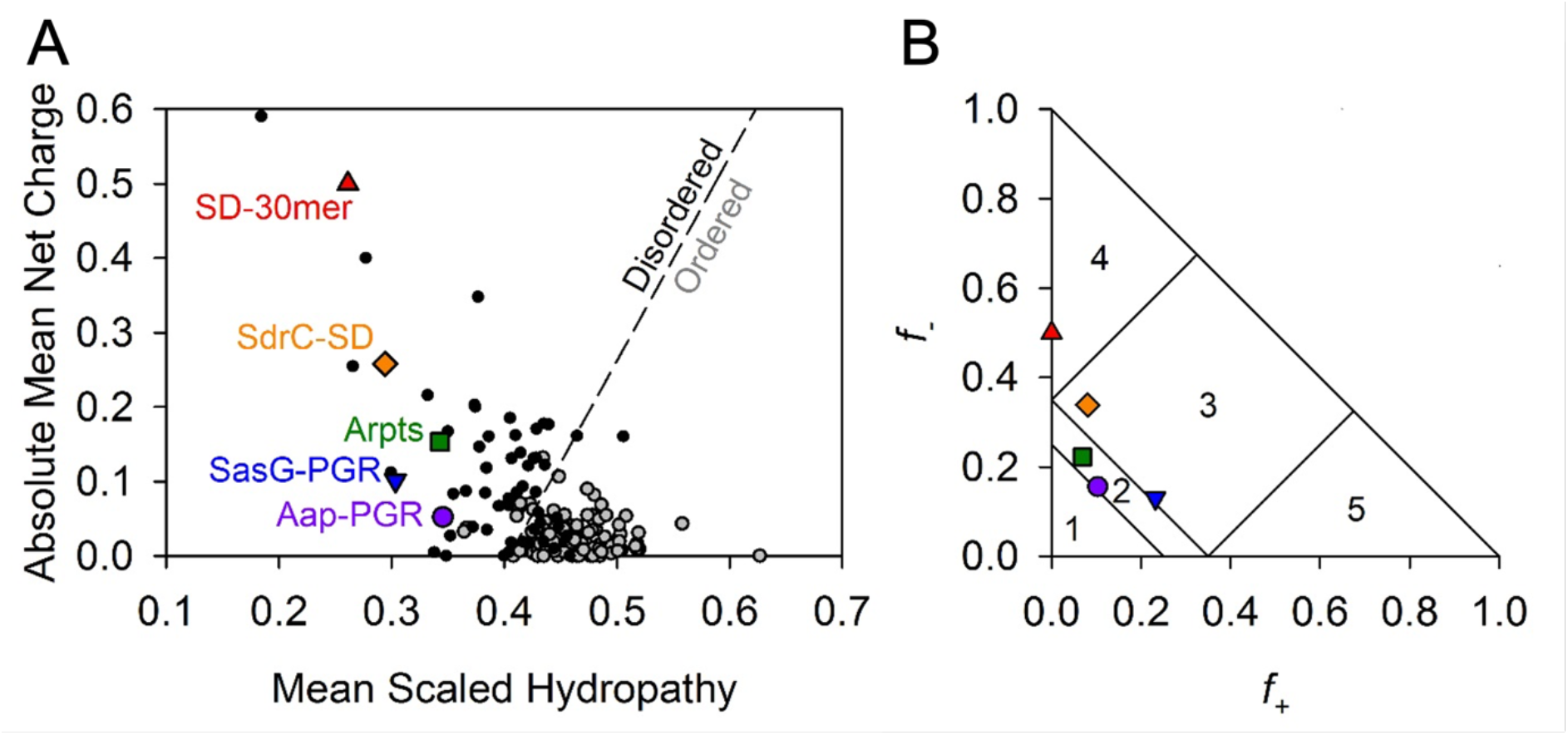
Predictions of disorder and classification of IDP constructs. The Uversky plot (A) is a powerful predictor of disorder based on net charge and hydrophathy [3]. A reference line (dashed) separates disordered proteins (black, filled circles) and ordered proteins (grey, filled circles). All IDP constructs of interest in this study fall on the “Disordered” side of the reference line. Panel (B) shows the IDPs plotted on the Das-Pappu phase plot [4]. Symbols are consistent with (A). Numbers refer to the phase plot region (See Table 1).

Several studies have been published examining the potential effects of charge on IDP conformation. Das & Pappu have investigated the impact of charge distribution or charge mixing on IDP conformational preferences [4]. They proposed classifying proteins based on the fraction of positively (*f*_+_) and negatively (*f*_-_) charged residues, resulting in the Das-Pappu plot shown in Figure 2B. Each region of this plot is discussed in detail elsewhere [4]. Region 2 contains Aap-PGR and Aap-Arpts. This region is considered a “boundary” region of so-called Janus sequences, where the conformational ensemble may sample either globule or tadpole conformations (Region 1) or coils, hairpins, or chimeras (Region 3). SdrC-SD and SasG-PGR both just cross the boundary into Region 3, populated by strong polyampholytes. For comparison in the Das-Pappu plot, we also included an SD-30mer comprised solely of repeating SD dipeptides, which is located in Region 4 of strong negatively charged polyelectrolytes that form swollen coils (Figure 2B). The difference between SdrC-SD and the SD-30mer is the inclusion of the native C-terminal 22 residues in SdrC that deviate from the SD repeating pattern. In addition to fractional charge, the specific distribution of charged residues is an important parameter that influences the configurational properties of polypeptide chains [4, 23]. Different parameters have been used to describe sequence charge distribution; both the κ parameter proposed by Das & Pappu [4] and sequence charge decoration (SCD) described by Sawle & Ghosh [23] report on the degree of charge clustering. Although they are calculated differently, a high value of κ or SCD represents highly segregated sequences (e.g., EEEEEKKKKK), which show a preference for hairpins or more compact conformations, while well-mixed sequences with low κ or SCD values (e.g., EKEKEKEKEK) experience self-avoidance due to electrostatic-repulsion and tend toward extended chains or Flory random coils [4, 23]. The κ and SCD values for each construct are shown in Table 1, along with additional parameters relevant to charge provided by the CIDER webserver [24].

**Table 1.** Sequence parameters for IDP constructs.

| Parameter | Aap-PGR | SasG-PGR | Aap-Arpts | SdrC-SD | SD-30mer |
| --- | --- | --- | --- | --- | --- |
| N | 135 | 69 | 189 | 62 | 30 |
| f- | 0.15556 | 0.13043 | 0.22222 | 0.33871 | 0.50000 |
| f+ | 0.1037 | 0.23188 | 0.06878 | 0.08065 | 0 |
| FCR | 0.25926 | 0.36232 | 0.29101 | 0.41935 | 0.50000 |
| NCPR | -0.05185 | 0.10145 | -0.15344 | -0.25806 | -0.50000 |
| Kappa | 0.05825 | 0.09562 | 0.08655 | 0.30207 | 0.02324 |
| SCD | 0.79 | 1.11 | 15.80 | 4.06 | 10.79 |
| FPR | 0.28889 | 0.17391 | 0.09524 | 0.03226 | 0 |
| Omega | 0.03234 | 0.07106 | 0.05146 | 0.00657 | 0.00096 |
| Hydropathy | 3.09259 | 2.72899 | 3.08466 | 2.64839 | 2.35 |
| Phase Plot Region | 2 | 3 | 2 | 3 | 4 |

The CIDER server [24] was used to calculate most parameters, including those required for the Das-Pappu Plot [4]; SCD was calculated as described [23].
**N:** Number of residues
**f-:** Fraction of negative residues
**f+:** Fraction of positive residues
**FCR:** Fraction of charged residues
**NCPR:** Net charge per residue
**Kappa:** $\kappa$ is a charge patterning parameter [4]. Highly mixed charged sequences approach $\kappa = 0$ , while highly segregated charged sequences approach $\kappa = 1$ .
**SCD:** Sequence charge decoration [23]. Well-mixed charged sequences approach $SCD = 0$ , whereas highly segregated charged sequences show large values of SCD.
**FPR:** Fraction of proline residues (not a parameter provided by CIDER, but included here for relevance)

A more relevant parameter in the case of Aap-PGR, SasG-PGR and Aap-Arpts might be omega, Ω. This value considers the patterning of proline and charged residues against all other residues [25]. While the implications of κ are dependent upon FCR, Ω is dependent on FCR and the number of proline residues. When used appropriately, such as a sequence with mostly similarly-charged charged residues and well-dispersed prolines, sequences with low Ω are more expanded or extended than those with high Ω [25]. One should use caution when the sequence of interest contains poorly mixed charged/proline residues, as this could enable long-range electrostatic attractions leading to hairpins, or there could be steric restrictions from groups of prolines. Table 1 also lists the Ω value for each construct. For Aap-PGR, SasG-PGR and Aap-Arpts, Ω is very low, suggesting the patterning of charged/proline residues could bias the conformational ensemble toward more extended conformations. SdrC-SD and SD-30mer contain few or no proline residues, so κ is the more appropriate parameter to consider for those constructs.

Uversky, et al. [3] and work from the Pappu Lab [4, 25, 27] have shown that charged residues can play an important role in the conformational preferences of IDPs, particularly when there is a high number of charged residues. Not surprisingly, there are additional factors that confer conformational bias. Work from the Whitten Lab has examined the effect of polyproline type-II (PPII) helix propensity [1, 5, 28] and α-helix propensity [29, 30] on the hydrodynamic radius (*R*_*h*_) of an IDP. PPII, thought to be a dominant backbone conformation in unfolded proteins [31], is a left-handed helix with three-fold rotational symmetry and is highly extended compared to an α-helix [32]. Tomasso, et al. [5] found the *R*_*h*_ values of IDPs could be predicted very well based on the PPII propensity and number of residues. Interestingly, the effect of charged residues was very weak compared to PPII propensity but could be significant where there was poor mixing of oppositely charged residues (i.e., sequences with a low κ value).

Parameters relevant to *R*_*h*_ prediction from PPII propensity are listed in Table 2. In the case of Aap-PGR, SasG-PGR and Aap-Arpts, there are a high number of prolines, which could place a strong PPII-based bias on their conformations. They are also rich in other residues that have a high propensity for PPII [5, 33-35]. The predicted fraction of residues in the PPII conformation, *f*_PPII_, is the averaged PPII propensity [5, 33] based on sequence composition.

**Table 2.**
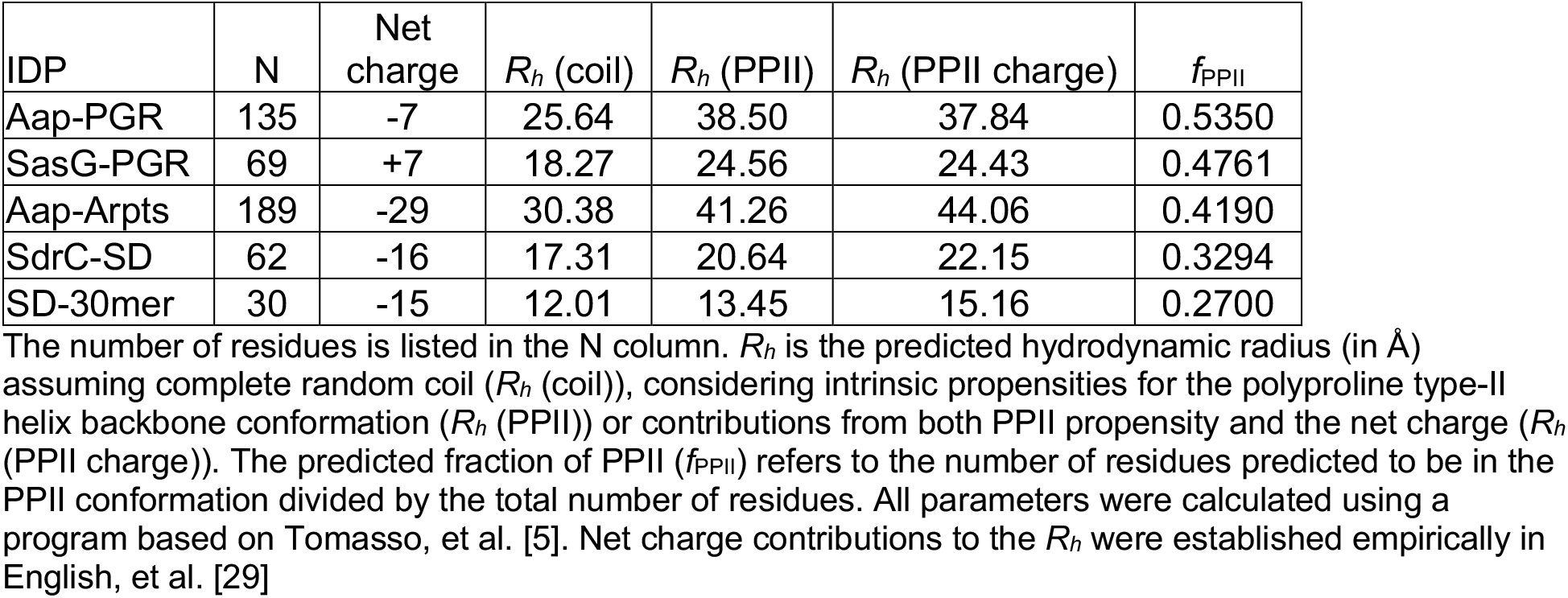
Calculated and predicted parameters of IDP constructs.

Aap-PGR has a higher *f*_PPII_ value than all other IDPs in the database used by Tomasso, et al. [5] (Table S1). SasG-PGR has the third highest *f*_PPII_ value, after the disordered tail of p53 (referred to as p53(1-93) Table S1) and Aap-PGR. Aap-Arpts lies at number six in the dataset of 27, while SdrC-SD and SD-30mer fall at or near the bottom of *f*_PPII_ values (Table S1). Also listed in Table 2 are the predicted *R*_*h*_ values assuming a random coil conformation that lacks strong preferences for backbone structures (R_h_ (coil)), predicted *R*_*h*_ for a random coil that samples PPII with a bias equivalent to *f*_PPII_ (R_h_ (PPII)), and predicted *R*_*h*_ for a random coil with a *f*_PPII_ bias and an additional effect owing to the net charge (R_h_ (PPII charge)). These values suggest which factors might influence the predicted *R*_*h*_ more strongly. Whether or not a protein is folded is perhaps the strongest influence on whether the global conformation is expected to be compact/globular or expanded/extended [1, 2, 5, 28, 36]. This *R*_*h*_ predictor assumes the sequence is unfolded. As expected, the contribution to the predicted *R*_*h*_ from PPII propensity is proportional to *f*_PPII_, and in the case of the IDPs with high FCR but low proline content, a contribution from the net charge can increase the predicted *R*_*h*_ significantly (Table 2).

### All IDPs are highly elongated monomers

Before examining the proteins by circular dichroism (CD), we examined their global conformations by sedimentation velocity analytical ultracentrifugation (AUC) and size-exclusion chromatography (SEC). Sedimentation velocity AUC measures the rate of sedimentation (sedimentation coefficient, *s*) as a species moves through solution [37, 38]. The calculated sedimentation coefficient (*s*) is dependent on the size (buoyant molar mass) and shape of the species. A protein that is highly expanded or elongated will sediment more slowly, due to increased frictional resistance, than a compact, globular protein of the same molar mass. The frictional coefficient is reported as the frictional ratio (*f*/*f*_0_), where the observed frictional coefficient is divided by the frictional coefficient of a sphere of the same volume [39]. We performed sedimentation velocity AUC on each of the IDPs of interest (for ease of comparison, previously published data for Aap-PGR [6] are included where appropriate alongside new data for SasG-PGR, Aap-Arpts, and SdrC-SD). The model-independent time-derivative analysis

(-dc/dt; Figure 3A) was performed for the datasets, with each distribution able to be fit with a single species of molar mass similar to the sequence-based monomer mass (Table 3). The c(*s*) distributions (Figure 3B) reveal a single boundary. Like with the dc/dt analysis, the c(*s*) analysis yielded estimated molar masses similar to monomer (see Table 3). In addition to determining the molar mass, the frictional ratio (*f*/*f*_0_) was also calculated and is listed in Table 3. In all cases, a very high *f*/*f*_0_ is observed, indicative of highly elongated global conformations. A slight overestimate of the molar mass was observed in all cases. This is likely due to the presence of hydrodynamic nonideality, which is expected to exist to some degree for such highly elongated proteins, despite the loading concentrations being at or below 1 mg/ml [40, 41].

**Table 3.** Sedimentation velocity AUC parameters.

| IDP | $M_{\text{seq}}$<br>(kDa) | $s_{20,w}$<br>(dc/dt) | $M_{\text{exp}}$<br>(kDa)<br>(dc/dt) | $s_{20,w}$<br>(c(s)) | $f/f_0$<br>(c(s)) | $M_{\text{exp}}$<br>(kDa)<br>(c(s)) | $R_s$<br>(Å) | $R_h$ (Å)<br>(SEC) | $R_h$ (PPII<br>charge) |
| --- | --- | --- | --- | --- | --- | --- | --- | --- | --- |
| Aap-PGR | 13.1 | 1.02 | 14.3 | 1.06 | 2.12 | 14.4 | 33.9 | 37.1 | 37.84 |
| SasG-PGR | 7.5 | 0.76 | 8.9 | 0.80 | 2.03 | 9.3 | 28.1 | 24.8 | 24.43 |
| Aap-Arpts | 19.1 | 1.25 | 25.6 | 1.30 | 2.52 | 22.2 | 46.2 | 40.8 | 44.06 |
| SdrC-SD | 6.4 | 0.91 | 8.4 | 0.97 | 1.93 | 8.1 | 24.7 | 21.1 | 22.15 |

**Figure 3.**
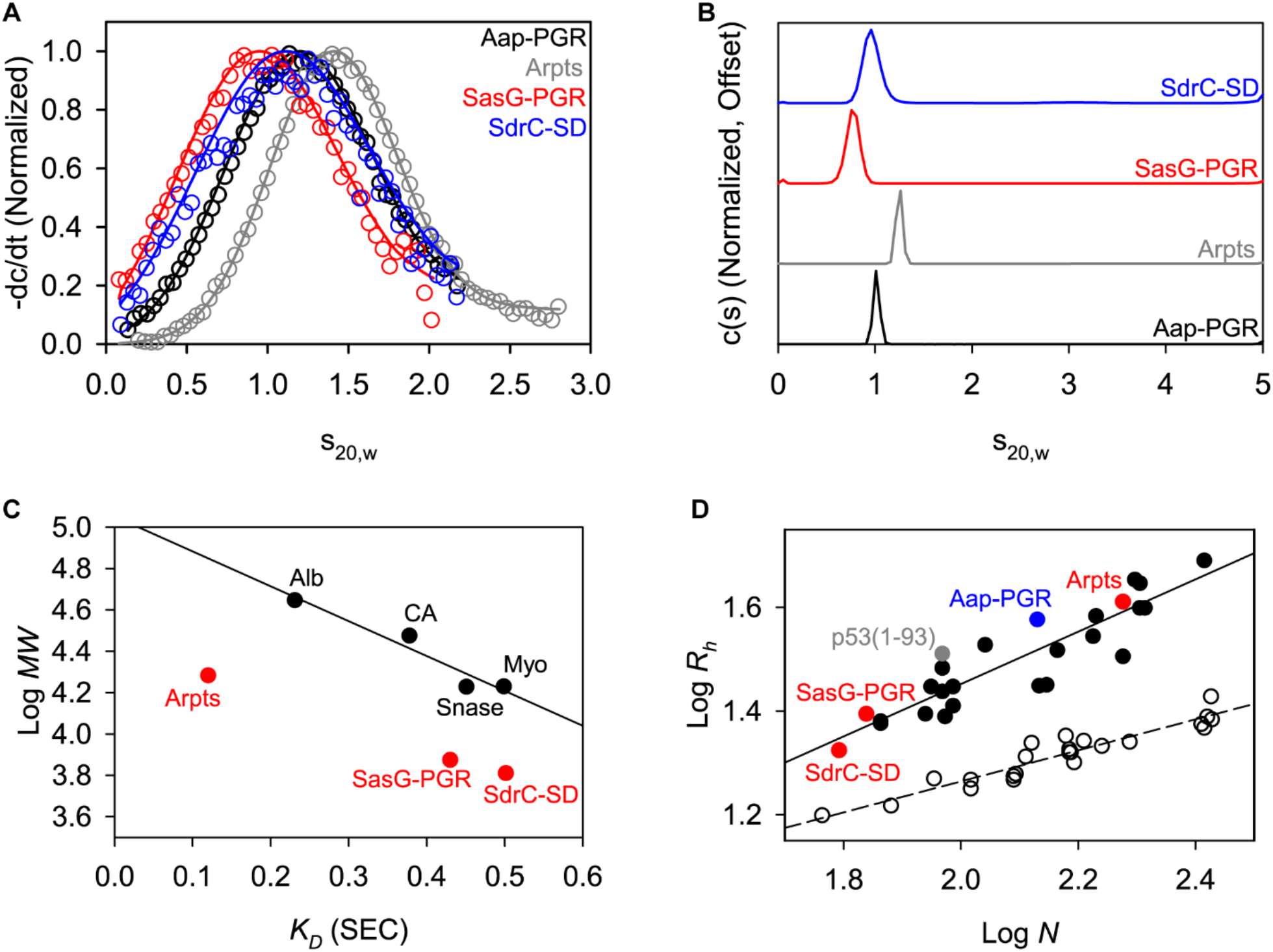
AUC and SEC indicate each construct is highly elongated and monomeric. A) Time-derivative distributions from sedimentation velocity data. Empty markers indicate data, with every 8^th^ data point shown for clarity. Solid lines represent the fit to a single-species model. B) Sedimentation velocity AUC data suggest each construct is monomeric and highly elongated, as indicated by the fitted parameters listed in Table 3. C) Linear relationship between SEC-measured K_D_ and logMW for the folded protein standards (Alb, albumin; CA, carbonic anhydrase; Myo, myoglobin; Snase, staphylococcal nuclease), showing that the IDP constructs all deviate from the trend. Aap-PGR is omitted from this plot, as it was previously analyzed using different column resin. D) Linear relationship between log N (number of residues) and logR_h_ in A. Folded proteins [1, 2] (open circles, dashed linear regression) and IDPs [5] (filled circles, solid linear regression) show distinct trends. SasG-PGR, Aap-Arpts, and SdrC-SD follow the trend with other IDPs, as previously observed for Aap-PGR [6].

In parallel to the AUC analysis, the hydrodynamic radius (*R*_*h*_) of each construct was determined experimentally by SEC as previously described [1, 6]. Each protein was run over a G-100 Sephadex column along with a panel of well-characterized globular control proteins that were used to determine the linear relationship between the thermodynamic retention factor (*K*_*D*_) and *R*_*h*_. We previously determined the *R*_*h*_ for Aap-PGR to be 37.06 Å, which was unusually high for a low MW protein of 13.1 kDa [6]. Similarly, the experimental R_h_ determination for SasG-PGR, Aap-Arpts, and SdrC-SD yielded relatively large values of 24.8, 40.8, and 21.1 Å, respectively (Figure 3C and Table 3). The correlation of predicted to experimental *R*_*h*_ was very good (R^2^=0.99; Figure S1). As with Aap-PGR, we observed that SasG-PGR, Aap-Arpts, and SdrC-SD cluster with other disordered proteins on a plot of log*R*_*h*_ vs log*N* (Figure 3D).

SEC-SAXS was used to additionally characterize the global conformation of the IDPs (Figure 4). The Guinier region of each dataset was linear and could be fitted to obtain an estimate of the radius of gyration (*R*_*g*_), as shown in Figure 4B and Table 4. The Kratky Plot (Figure 4C) indicates extended particle shape due to the upward trend in the data as *qR*_*g*_ increases. The Kratky indications are confirmed by the P(r) distributions shown in Figure 4D, which indicate elongated shape by the tailing of the distribution, in contrast to the Gaussian shape expected for a spherical particle. Using the ratio, *ρ*, of the SAXS-measured *R*_*g*_ and SEC-measured hydrodynamic radius (*R*_*h*_), another estimate of the shape can be obtained. A globular protein is expected to yield a *ρ* of ∼0.7, while higher values indicate elongation [42, 43]. All constructs analyzed yielded values of *ρ* ranging from 0.98 to 1.17, consistent with an elongated state in solution (Table 4).

**Table 4.** SEC-SAXS Analysis Results.

| IDP | $R_g$ (Å)<br>(Guinier) | $R_g$ (Å)<br>(P(r)) | $D_{max}$ (Å)<br>(P(r)) | $R_h$ (Å)<br>(SEC) | $\rho$ ($R_g/R_h$) |
| --- | --- | --- | --- | --- | --- |
| Aap-PGR | $35.0 \pm 0.3$ | $36.4 \pm 0.2$ | 150 | 37.1 | 0.98 |
| SasG-PGR | $25.5 \pm 0.2$ | $26.6 \pm 0.1$ | 108 | 24.8 | 1.07 |
| Aap-Arpts | $46.6 \pm 0.5$ | $47.8 \pm 0.2$ | 182 | 40.8 | 1.17 |
| SdrC-SD | $23.4 \pm 0.4$ | $24.2 \pm 0.4$ | 104 | 21.1 | 1.15 |

**Figure 4.**
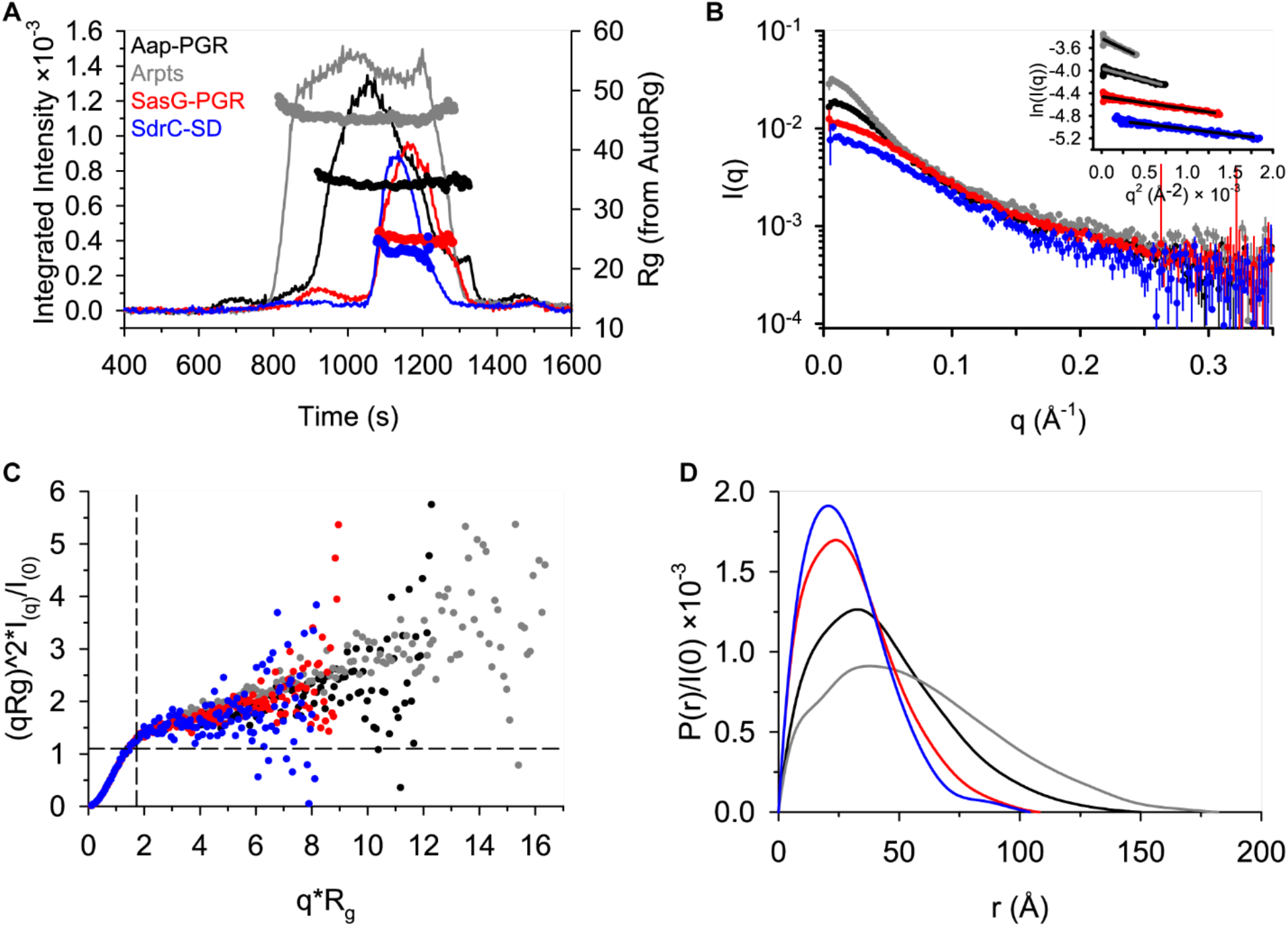
SAXS data confirm highly extended conformations for all constructs. A) SEC-SAXS elution profiles showing scattering intensity (after buffer subtraction) and *R*_*g*_ calculated using AutoRg (markers). B) The scattering profiles for each dataset, following the same color scheme as (A). The inset shows the Guinier range with lines representing the Guinier fit. C) The normalized Kratky plot. A spherical particle should exhibit a Gaussian shaped distribution with a maximum at the intersection of the dashed lines. D) The *P(r)* distributions normalized by *I(0)*. The *D*_*max*_ is the maximum radius in the distribution.

**Figure 4.**
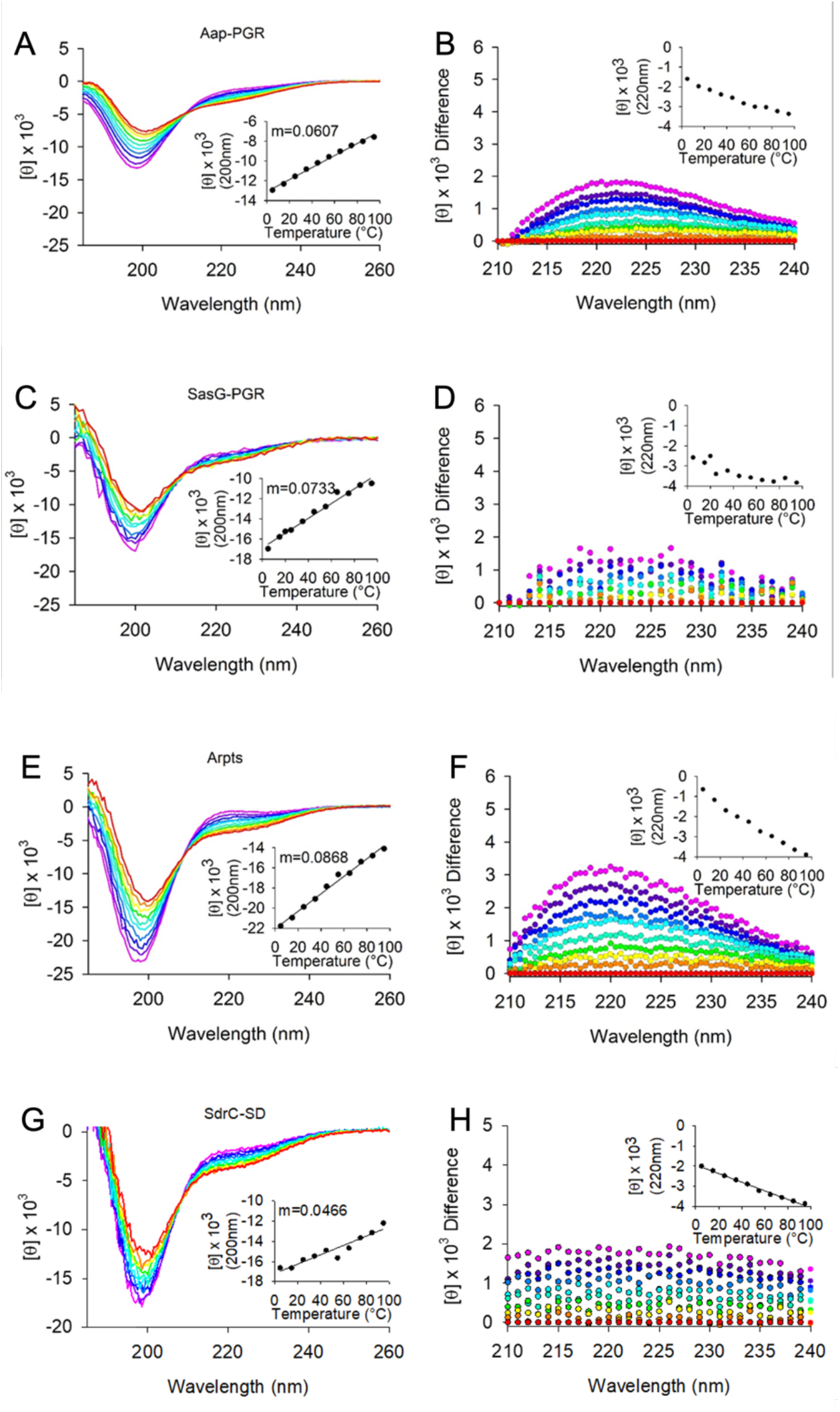
Circular dichroism wavelength scans show constructs have primarily random coil and PPII helix content. (A) shows the wavelength scans for Aap-PGR from 5°C-95°C (cool to warm colors), while (B) shows the difference of each scan minus the 95°C scan. This highlights the shift in the local maximum around 220 nm. The insets highlight the linearity of the CD signal at 200 nm (A) or 220 nm (B). Data are shown for SasG-PGR (C, D), Aap-Arpts (E, F), and SdrC-SD (G, H). The slope determined by linear regression of the CD signal (200 nm) vs temperature is listed as “m.”

### CD confirms random coils with PPII bias

After confirming the constructs of interest are monomeric and highly extended, we utilized circular dichroism (CD) to measure the secondary structure of the IDPs. Random coil exhibits a minimum near 200 nm, while PPII exhibits an even stronger minimum near 200 nm, along with a weak local maximum near 220 nm [44]. It is very useful to measure the temperature dependence of the CD spectrum to examine the transition to random coil at high temperatures. PPII is stabilized at low temperatures, whereas high temperatures disrupt PPII, resulting in random coil [6, 28, 44, 45]. The resulting temperature dependence is usually linear, which indicates little or no cooperativity in PPII formation [6, 45, 46]. However, if there is significant α-helix or β-sheet content, one would expect to observe a sigmoidal response in the temperature dependence [47].

Figure 4 shows CD wavelength scans of Aap-PGR, SasG-PGR, SdrC-SD, and Aap-Arpts as a function of temperature. In each case, it is apparent that the primary features of the scans are the 200 nm minimum and the shift in intensity of the 220 nm signal. This strongly indicates that each IDP is composed of a mix of random coil and PPII. The insets of Figure 4A, C, E, and G show the linear trend of the CD signal at the 200 nm minimum. For Aap-PGR, SasG-PGR, and Aap-Arpts, the slope (*m*) of the linear regression of these data trends inversely with *f*_PPII_, where higher PPII propensity was associated with a shallower slope.

### Cosolvents perturb secondary structure to varied degrees

In Figure 5, we examined the effect of denaturants on the CD signal. Urea and guanidinium hydrochloride (GdnHCl) destabilize the folded state of globular proteins via favorable interactions with the peptide backbone and hydrophobic residues [48]. Generally, GdnHCl is at least twice as effective as urea at denaturing folded proteins, but differences can be expected depending on amino acid composition. Interestingly, once a protein is unfolded, these denaturants will induce PPII [46, 49-51]. Figure 5 focuses on the region around the local maximum at 220 nm. The strong absorbance of urea and GdnHCl preclude reliable measurements below this wavelength range. Aap-PGR showed only a slight increase in PPII, indicated by the lack of significant change in the CD signal at 220 nm in the presence of urea or GdnHCl. SasG-PGR, Aap-Arpts, and SdrC-SD showed larger increases around 220 nm, especially at 4°C where PPII is more stable, suggesting a larger increase in the PPII content upon urea and GdnHCl addition. As seen with the temperature dependence in Figure 4, the magnitude of the changes is (qualitatively) inversely proportional to *f*_PPII_ for Aap-PGR, SasG-PGR, and Aap-Arpts, but the data for SdrC-SD most closely resemble Aap-PGR despite having a low *f*_PPII_ and higher values of κ and SCD. A possible interpretation of these data is that Aap-PGR does not show much of an increase in PPII content, because it already has a high PPII content (high *f*_PPII_), whereas Aap-Arpts has a lower inherent PPII content in its native state and thus is more susceptible to the further denaturant-induced PPII transition. Figure 6 shows similar CD measurements in the presence of a stabilizing osmolyte, TMAO, or an alcohol, TFE. TMAO is a cosolvent that is capable of inducing compact conformations (often the functional native state for globular folds), primarily through strongly unfavorable interactions with the peptide backbone. TMAO is particularly useful, because it does not force a specific structure, it simply destabilizes the unfolded or expanded state, which results in a shift in the equilibrium from unfolded states to the folded, native state [48, 52-54]. This is in stark contrast to TFE, which induces α-helix formation, even when this is non-native [50, 55-58].

**Figure 5.**
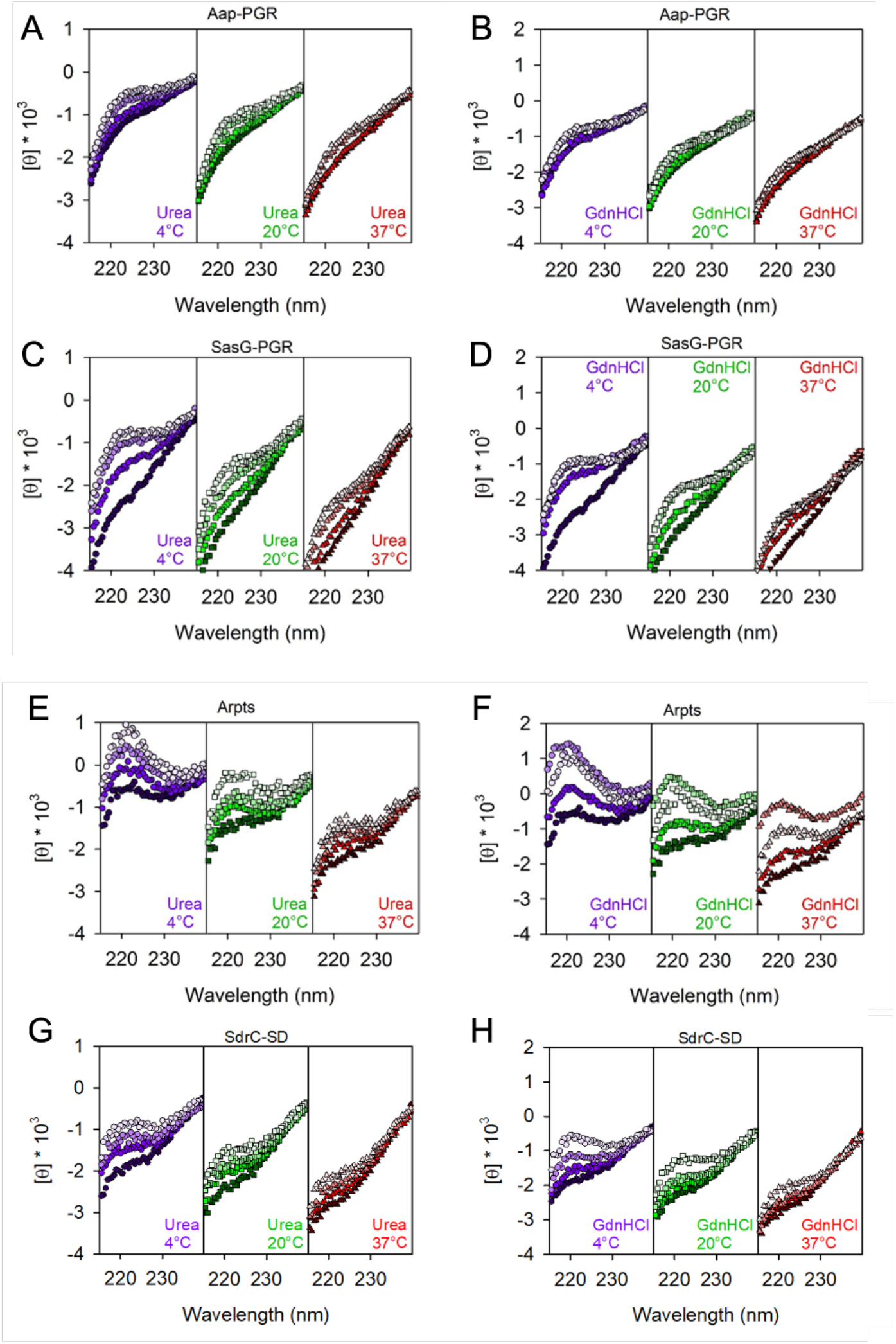
Constructs respond to denaturants to different extents. Concentrations of urea were 0, 2, 4, 6 M, colored from darkest to lightest fill. Plots show 0, 2, 4, 6 M GdnHCl using the same color scheme.

**Figure 6.**
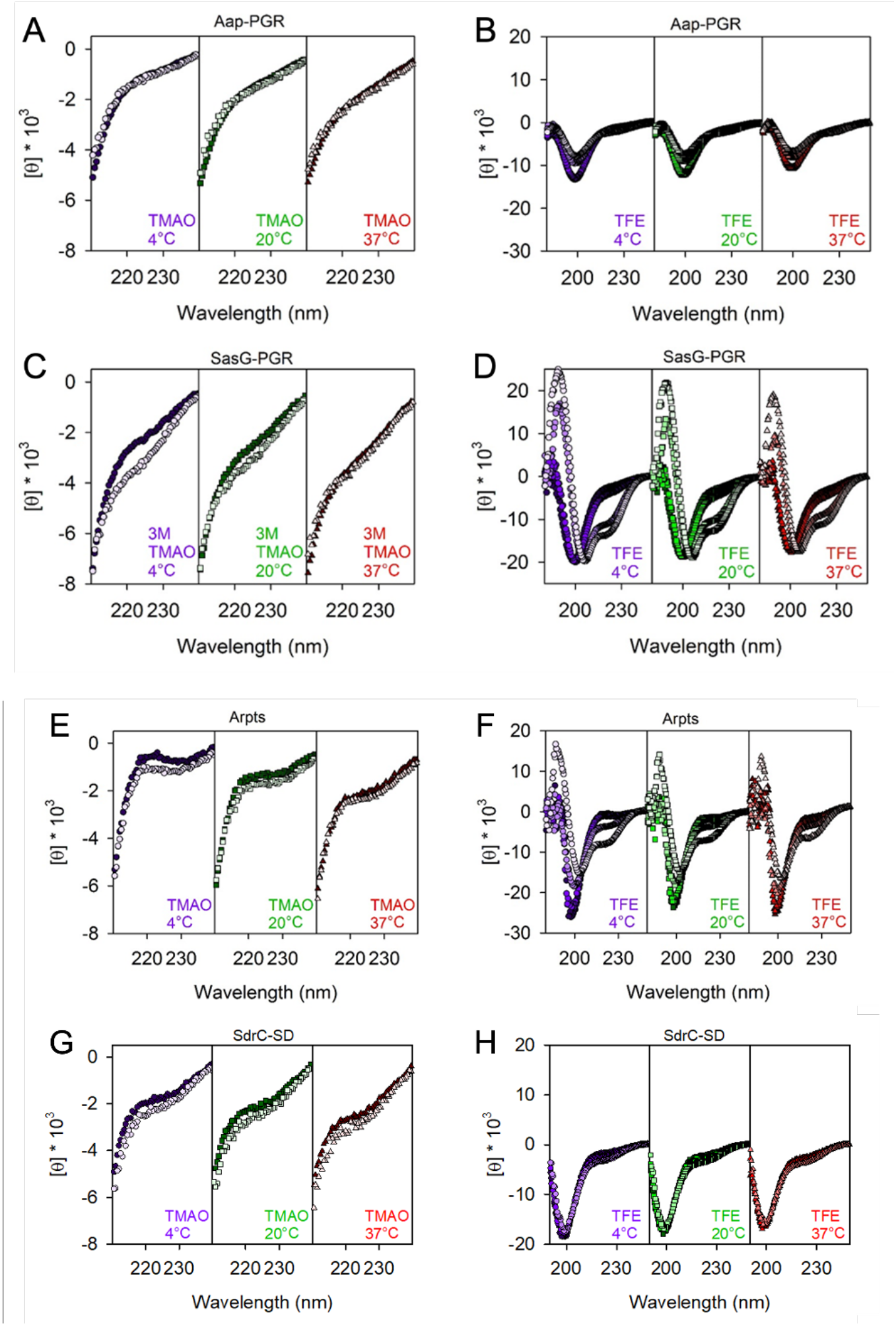
Comparison of IDP responses to TMAO and TFE. Concentrations of TMAO (A, C, E, and G) were 0 (dark fill) and 3 M (light fill). The ability for TFE to perturb the secondary structure is shown in (B, D, F, and H). TFE concentrations were 0, 15%, 45% and 75% (from dark fill to light fill) except for panel H, where 75% TFE led to aggregation of SdrC-SD.

The addition of TMAO to Aap-PGR or SdrC-SD has essentially no effect (Figure 6A, 6D). We previously hypothesized this was due to the moderately high frequency of charged residues, which form favorable interactions with TMAO, and therefore, are seemingly able to counter the backbone-TMAO effects [6, 52]. If this is a valid explanation for the lack of effect, then we should also see a similar result for SasG-PGR and Aap-Arpts, which have a greater fraction of charged residues (Table 1). Indeed, we see a little to no effect of TMAO on the CD spectrum of these two IDPs, although some difference is observed at low temperatures in both cases (Figure 6). A lack of response to TMAO has been observed in other IDPs, including myelin basic protein [59] and Starmaker-like protein [60], which also have high FCR values of 24% and 47%. These two proteins have very low proline content (3% and 6%), so while proline also has a favorable interaction with TMAO, the charged residues are likely the most dominant factor in these cases [52]. However, the decreased proline content in SasG-PGR and Aap-Arpts (Table 1) may allow for the slight effect of TMAO (Figure 6). The lack of a response to TMAO could also suggest that the native state is already populated, and there is not a separate, compact state which might be induced upon binding of a ligand as is the case with some IDPs [61, 62].

The response to TFE was much greater than TMAO (Figure 6). As discussed elsewhere, Aap-PGR has an apparent shift from PPII to random coil upon TFE addition, but no indication of α-helix formation [6]. This is not surprising due to the high frequency of proline, which sterically prohibits α-helix formation. However, we expect that with the lower frequency of proline in SasG-PGR and Aap-Arpts, TFE might have a stronger effect. Indeed, we do see a much more significant effect of TFE on SasG-PGR and Aap-Arpts. CD spectra of α-helix content shows minima at 222 nm and 208 nm [47]. Interestingly, while both of these proteins show clear development of the local minimum around 222 nm and a shift of the ∼200 nm minimum toward the 208 nm minimum, there are potentially interesting differences. For SasG-PGR, there is little or no weakening of the ∼200 - 208 nm minimum, while Aap-Arpts (and Aap-PGR) show a strong weakening of this signal. Once again, the TFE response seen for SdrC-SD shows striking resemblance to that of Aap-PGR, despite the differences in f_PPII_, f-, FCR, NCPR, and κ or SCD for the two sequences. Despite the paucity of prolines in SdrC-SD, this highly negatively charged construct showed almost a complete resistance to TFE-induced folding into α-helical secondary structures.

## Discussion

After our investigation of Aap-PGR, the proline/glycine-rich region from Aap, we became curious if other low-complexity, stalk-like regions would show similar extended conformations and resistance toward compaction [6]. Therefore, we chose several additional regions to investigate and compare to Aap-PGR. These regions include the similar stalk-like region of SasG, the *S. aureus* ortholog of Aap (*S. epidermidis*), and the serine-aspartate region of SdrC, which spans the space between the functional domain(s) and the cell wall-anchoring motif of this protein [20] The Arpt region of Aap was included as a control IDP sequence that does not function as a stalk.

We began by showing sequence-based predictions suggesting these proteins will be disordered in solution. The amino acid compositions alone, which lacked many hydrophobic residues (order-promoting residues) and were enriched with polar and charged residues (disorder-promoting residues), are strong indicators of disorder [61, 63]. Furthermore, our sequences of interest have a wide range of net charge (Figure 2, Table 1). This allowed for evaluation of potential charge effects on conformation. We also predicted the hydrodynamic radius (*R*_*h*_) of each sequence based on PPII propensity and charge effects, which suggested PPII was a strong contributor to *R*_*h*_ in sequences with medium or high proline content, but charge effects might be much more dominant in the sequences lacking prolines. Sedimentation velocity AUC showed that each IDP is monomeric and exists in highly elongated conformations (Figure 3 and Table 3). The calculated radius of gyration (*R*_*g*_) and Stokes-Einstein radius (*R*_*s*_) trends well with both the experimentally determined *R*_*h*_ and the predicted *R*_*h*_ based on PPII and charge (Table 3).

Using circular dichroism (CD), we verified that Aap-PGR, SasG-PGR, Aap-Arpts, and SdrC-SD exist in a PPII-random coil equilibrium, which can be modulated by temperature (Figure 4). The dependence of the CD spectra on temperature correlated with the predicted *f*_PPII_ for the sequences with low values of kappa (i.e., Aap-PGR, SasG-PGR, and Aap-Arpts), whereas SdrC-SD behaved like Aap-PGR despite a low predicted *f*_PPII_. Likewise, the ability of chemical denaturants to increase the PPII signal showed an inverse trend with *f*_PPII_ for the low-kappa constructs; specifically, there was a notable increase in PPII with denaturant concentration in the sequences with lower PPII propensity, except for SdrC-SD which showed moderate PPII increases with denaturant concentration.

With all IDPs tested, there was very little change in the CD spectra upon TMAO addition (Figure 6). This is likely due to the high fraction of charged residues, with some additional contribution from proline content. It may also suggest that these IDPs do not form additional compact states, but actually remain disordered in their native states. This is in line with our hypothesis stating that SasG-PGR and SdrC-SD both form an extended stalk that is resistant to compaction, like Aap-PGR [6]. The addition of α-helix-inducing TFE had more significant effects than TMAO on all low-kappa IDPs (except SdrC-SD), but development of the characteristic 208 and 222 nm α-helix minima was very clear in particular for SasG-PGR and Aap-Arpts. In Aap-PGR, there are frequent prolines in every third position, making it more unlikely that helices could form in these regions. SasG-PGR also contains a proline in every third position throughout the first half of the region, however, the second half becomes more variable and lacks proline residues. This likely allows significant α-helix formation to occur in the latter half of the SasG-PGR sequence in the presence of TFE. While Aap-Arpts sequence does contain a fair number of prolines, they are more spaced out (∼11-15 residues apart) which probably allows for short, interspersed α-helices to form (the average length of an α-helix is 10-15 residues [64]).

Notably, the Aap-Arpts sequence is distinct from the other constructs tested in that its location at the N-terminus of Aap is inconsistent with a stalk-like function; although it showed minimal change in its far-UV CD spectrum in the presence of TMAO, it was able to form a significant degree of helical content at high TFE concentrations. The role of the A-repeat region of Aap is unclear, but its TFE response indicates that Aap-Arpts could undergo induced folding, for example upon ligand binding.

Remarkably, Aap-PGR and SdrC-SD, the two stalk constructs that were most divergent in sequence pattern, showed nearly identical behavior in solution. Both formed highly extended conformations by AUC and SAXS, demonstrated the least temperature dependence (based on the inset CD slopes in Figure 4), and showed minimal response to denaturants or osmolytes. In particular, both Aap-PGR and SdrC-SD were highly resistant to compaction, based on their nearly complete lack of response to TFE or TMAO. Despite these functional similarities, Aap-PGR and SdrC-SD occupy disparate regions of the Das-Pappu plot, with Aap-PGR falling in the boundary region of Janus sequences and SdrC-SD occupying the strong polyampholyte region. Among the sequences tested, Aap-PGR and SdrC-SD are on opposite extremes in terms of FCR, FPR, and kappa; the only similarity based on sequence parameters is Ω, since both sequences have lower values compared to SasG-PGR or Aap-Arpts. Thus, our data indicate that both high proline/PPII content (in Aap-PGR) and high electrostatic repulsion between negative residues (in SdrC-SD) result in similar self-avoiding configurations that result in highly extended, compaction-resistant stalk-like behavior.

It is very common for staphylococcal CWA proteins to feature low-complexity sequences predicted to be intrinsically disordered at the extreme C-terminus; these regions presumably function as an extended stalk. Table 5 lists a series of CWA proteins containing C-terminal LPXTG motifs from *S. aureus* or *S. epidermidis* with known or suspected adhesion functions and describes the characteristics of the low-complexity sequence regions upstream of the LPXTG sortase motifs. Of these 17 CWA proteins with adhesin-like functions, 82% have either SD-rich repeats (9 of 17) or proline-rich repeats (5 of 17) at the C-terminus that are expected to show highly elongated, compaction-resistant stalk behavior as seen for SdrC-SD and Aap-PGR. The results from this work indicate that such stalk regions share functional similarity that can be encoded by highly divergent sequence patterns such as high PPII propensity due to repeating proline residues or electrostatic repulsion due to charge effects (e.g., high values of f- or kappa). The necessity for multiple mechanisms to achieve similar stalk-like behavior might be related to environmental conditions in distinct niches (e.g., colonizing skin versus forming biofilm on heart valves or an implanted device) or to specific interactions with other macromolecules needed for particular functions. For example, the stalk region may engage with various macromolecular components comprising the biofilm matrix. A stalk with a high charge density (e.g. SD repeats in the Sdr family of proteins) could interact with the positively-charged biofilm polysaccharide PIA/PNAG or with zwitterionic teichoic acids in the biofilm matrix. Furthermore, context-dependent modification of particular sequences in stalk regions can also provide functional specialization; for example, glycosylation of SD repeats of ClfA, ClfB, SdrC, SdrD, and SdrE by specific glycosyltransferases confers resistance to proteolysis [65] and glycosylation of the SD-rich region of Pls promotes biofilm formation [66]. Approaches such as swapping stalk regions between cell wall-anchored proteins among different bacteria or reducing the PPII propensity of specific stalk regions and testing the capacity for biofilm formation or host tissue adherence would provide additional insights into the biological implications of these observations.

**Table 5.** Stalk regions in adhesin-like staphylococcal CWA proteins.

| Protein | Ligand(s) | Length:<br>LCR<br>(Total) | SD<br>repeats | Prolines | Das-Pappu<br>region | FCR | FPR | $f_{PPII}$ | NCPR | Kappa | SCD | Omega |
| --- | --- | --- | --- | --- | --- | --- | --- | --- | --- | --- | --- | --- |
| **SD-rich** |  |  |  |  |  |  |  |  |  |  |  |  |
| SdrC | Self,<br>β-neurexin | 170<br>(193) | 83 | 2 | 4 | 0.505 | 0.010 | 0.289 | -0.479 | 0.056 | 110.90 | 0.00195 |
| SdrD | Desmoglein-1 | 142<br>(165) | 71 | 2 | 4 | 0.466 | 0.012 | 0.290 | -0.415 | 0.107 | 82.04 | 0.00348 |
| SdrE | Factor H | 158<br>(181) | 79 | 2 | 4 | 0.467 | 0.011 | 0.288 | -0.406 | 0.125 | 99.61 | 0.00299 |
| SdrF<br>(*S. epi*) | Collagen | 170<br>(193) | 84 | 2 | 4 | 0.470 | 0.010 | 0.289 | -0.414 | 0.114 | 113.55 | 0.00280 |
| SdrG<br>(*S. epi*) | Fibrinogen ($\beta$) | 140<br>(163) | 69 | 2 | 4 | 0.472 | 0.012 | 0.293 | -0.420 | 0.106 | 79.96 | 0.00264 |
| Pls | Unknown | 294<br>(309) | 112 | 1 | 4 | 0.466 | 0.003 | 0.286 | -0.405 | 0.125 | 325.06 | 0.00298 |
| ClfA | Fibrinogen ($\gamma$),<br>Factor I | 323<br>(352) | 130 | 10 | 4 | 0.423 | 0.028 | 0.289 | -0.401 | 0.101 | 260.80 | 0.03617 |
| ClfB | Fibrinogen ($\alpha$),<br>Cytokeratin 8,<br>Cytokeratin 10,<br>Loricrin | 252<br>(286) | 99 | 20 | 4 | 0.451 | 0.070 | 0.337 | -0.409 | 0.105 | 196.05 | 0.04653 |
| SesJ<br>(*S. epi*) | Plasminogen | 213<br>(217) | 36 | 0 | 4 | 0.378 | 0 | 0.288 | -0.378 | 0.052 | 121.12 | 0.02044 |
| **Pro-rich** |  |  |  |  |  |  |  |  |  |  |  |  |
| Aap<br>(*S. epi*) | Self | 126<br>(138) | 0 | 40 | 2 | 0.261 | 0.290 | 0.537 | -0.043 | 0.058 | 0.40 | 0.03154 |
| SasG | Self | 40<br>(69) | 0 | 12 | 3 | 0.362 | 0.174 | 0.476 | 0.101 | 0.096 | 1.11 | 0.07106 |
| CNA | Collagen,<br>C1q | 55<br>(58) | 0 | 15 | 3 | 0.431 | 0.259 | 0.554 | 0.052 | 0.071 | -0.08 | 0.09578 |
| FnbpA | Fibronectin,<br>Fibrinogen ($\gamma$) | 76 (108) | 0 | 36 | 2 | 0.269 | 0.333 | 0.596 | -0.046 | 0.209 | -1.76 | 0.04935 |
| FnbpB | Fibronectin,<br>Fibrinogen ($\alpha$),<br>Elastin | 62<br>(94) | 0 | 29 | 2 | 0.277 | 0.309 | 0.584 | -0.021 | 0.205 | -1.74 | 0.05606 |
| **Other** |  |  |  |  |  |  |  |  |  |  |  |  |
| SraP<br>(SasA) | Sialoprotein,<br>gp340 | 73<br>(80) | 2 | 2 | 1 | 0.138 | 0.025 | 0.318 | -0.037 | 0.153 | -0.06 | 0.45721 |
| FmtB<br>(SasB) | Unknown | 57<br>(58) | 0 | 1 | 1 | 0.190 | 0.017 | 0.368 | 0.052 | 0.342 | -0.40 | 0.28539 |
| SasC | Self | 177<br>(220) | 2 | 9 | 2 | 0.323 | 0.041 | 0.390 | -0.023 | 0.147 | -0.06 | 0.12394 |

Ligand specificities for the LPXTG proteins are according to [22, 67-71]; sequence parameters were calculated using the CIDER server (http://pappulab.wustl.edu/CIDER/) [4, 25], $f_{PPII}$ was calculated using the Tomasso et al. algorithm [5], and SCD was calculated using the equation for SCD provided in Sawle & Ghosh [23].

## Materials and methods

### Cloning and protein expression

SasG-PGR, Aap-Arpts and SdrC-SD genes were ordered as IDT gBlocks with 5’ CACC sequences for cloning into the Gateway Cloning System via pENTR/D-TOPO reaction, followed by TEV cleavage sites (ENLYQF/G) and the sequences in Figure 1, followed by a stop codon. The gene was transferred from the pENTR vector to a destination vector containing an N-terminal His_6_ tag and maltose binding protein (MBP), which was kindly provided by Dr. Artem Evdokimov. Aap-PGR DNA was synthesized by LifeTechnologies GeneArt® as previously described and moved into the same His_6_-MBP destination vector [6]. SD-30mer was ordered as a peptide from Peptide 2.0 at ≥95% purity.

BLR(DE3) cells were transformed with the destination vector containing the gene of interest. Overnight cultures were grown at 37 °C, then 25 ml used to inoculate 1 l LB broth containing ampicillin and tetracycline antibiotic selection. The cultures were grown to an OD_600_ of 0.8 - 1.0 at 37 °C, shaking at 200 - 250 rpm. The cultures were then placed in an ice bath until cooled to 10 °C. At this point, ethanol was added to 2% final concentration (volume/volume) and IPTG to 200 µM. The cultures were placed back into a shaker to incubate overnight at 20 °C. The following morning, cultures were centrifuged at 4,500 rpm for 1 hr, the supernatants discarded, and the pellets resuspended in 20 mM Tris (pH 7.4) and 300 - 500 mM NaCl. Resuspended pellets were stored at -20 °C.

### Protein purification

Frozen pellets were thawed, sonicated to lyse the bacteria, centrifuged for 45 min at 14,000 rpm, the supernatant filtered through a 0.22 µm filter, and the protein purified by Ni^2+^-affinity chromatography using an Äkta FPLC Chromatography System (GE Healthcare). The fusion proteins were eluted with a gradient of 1 M imidazole and dialyzed into 20 mM Tris (pH 7.4) and 300 mM NaCl. The protein was then incubated with TEV protease for 6 - 8 hours at room temperature. A subtractive Ni^2+^-affinity step was performed to capture uncleaved His_6_- tagged fusion protein, His_6_-tagged MBP, and His_6_-tagged TEV protease, while cleaved protein of interest did not interact with the Ni^2+^-affinity column. The cleaved protein of interest was then purified by size exclusion chromatography using a Superdex 75 prep grade column (GE Healthcare). Where necessary, only fractions containing the highest purity of full-length protein was collected from the Superdex 75 elution, or ion exchange chromatography was used to remove contaminants or degraded species.

### Analytical ultracentrifugation

Sedimentation velocity experiments were performed on a Beckman Coulter XL-I AUC, using 1.2 cm two-sector epon-charcoal centerpieces. Experiments were performed at 48,000 rpm at 20 °C in a An-60 Ti rotor and were continued until sedimentation was complete or back-diffusion became obvious in the raw data (∼20-24 hours). Interference optics were used alongside absorbance near 230 nm. Concentration estimates were obtained from the interference data. The samples were examined after dialysis into 20 mM KPO_4_, 150 mM NaCl, pH 7.4 or immediately following size exclusion chromatography. Data analysis was completed in SEDFIT using the continuous c(s) distribution model [39] and DCDT+ [72, 73] with fitting performed against the dc/dt distribution [74] using a single species model. SEDNTERP was used to estimate partial specific volume, buffer density and viscosity, and the fringes per mg/mL at 655 nm [75].

### Circular dichroism

An Aviv 215 spectrophotometer was used to perform CD measurements. The instrument is equipped with an Aviv peltier junction temperature control system. A 0.05 cm quartz cuvette (Hellma Analytics) was used for all measurements. Temperature-dependence experiments were performed using protein dialyzed into 20 mM potassium phosphate (pH 7.4) and 50 mM NaF. For cosolvent experiments, concentrated protein samples were mixed with appropriate amounts of cosolvent or water to match protein concentrations within experiments. Urea used was 8 M ultra-pure grade solution (Amresco), GdnHCl was 8 M high purity solution (Pierce), TFE was >99.0% purity (Sigma-Aldrich), and TMAO was 95% solid (Sigma-Aldrich) which was dissolved into water before addition to samples. All protein-cosolvent spectra were subtracted from buffer blanks which contained equal amounts of cosolvent at each temperature.

To convert data to mean residue ellipticity, [θ], Equation 1 was used. In this equation, θ is raw machine units output by the instrument, MRW is the mean residue weight, l is the path length of 0.05 cm, and c is the concentration in mg/ml units.

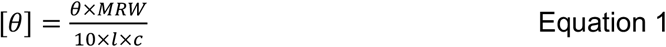

### Small-angle X-ray scattering

SAXS data were collected at BioCAT (beamline 18ID at the Advanced Photon Source, Chicago, Illinois, US) with in-line size-exclusion chromatography (SEC). Samples were dialyzed into 20 mM KPO_4_, 150 mM NaCl, pH 7.4. A Superdex 75 Increase 10/300 GL column (Cytiva) was used in conjunction with an Äkta Pure FPLC (GE). The flow rate was 0.6 mL/min, the elution was monitored by UV at 280 nm, and the flow path was directed through the SAXS flow cell. The flow cell consisted of a 1 mm (internal diameter) quartz capillary with ∼20 µm walls. A coflowing buffer sheath separated sample from capillary walls, helping to prevent radiation damage [76].

Scattering intensity was recorded using a Pilatus3 × 1M (Dectris) detector which was 3.6 m from the sample, yielding a q-range of 0.005 Å^-1^ to 0.35 Å^-1^. Exposures of 0.5 s were acquired every 1 s during elution. Data reduction was performed using BioXTAS RAW 1.6.0 [77]. Buffer blanks were created using regions flanking the elution peaks which showed baseline scattering intensity. I(q) vs q curves were produced by subtracting buffer blanks from elution peaks and used for downstream analysis. SAXS analysis was performed using BioXTAS RAW 2.1.0 [77], which called various functions from ATSAS 3.0.3 [78, 79].

### Size-exclusion chromatography

Sephadex G-100 (GE Healthcare) was equilibrated in 10 mM sodium phosphate (pH 7.0) and 100 mM NaCl. A Bio-Rad BioLogic LP System was used to monitor UV absorbance at 280 nm to determine elution volumes (*V*_*e*_). Concentrated samples of SasG-PGR, Aap-Arpts, and SdrC-SD at 2-3 mg/ml and at a volume of 80 µL were measured by SEC, as previously described for Aap-PGR [6]. To determine the void (*V*_*0*_) and total column volume (*V*_*t*_), we ran 10 μL of 3 mg/ml blue dextran and 0.03 mg/ml 2,4-dinitrophenyl-L-aspartate through the column separately from the proteins. The thermodynamic retention factor *K*_*D*_ was calculated using Eq. (2):

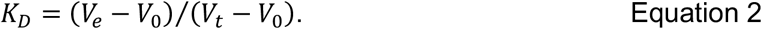

To determine the hydrodynamic radius of SasG-PGR, Aap-Arpts, and SdrC-SD, *R*_*h*_ measured previously by dynamic light scattering (DLS) for a set of globular protein standards were plotted against the experimentally determined *K*_*D*_ values. The protein standards were nuclease, carbonic anhydrase, myoglobin and albumin; DLS-measured *R*_*h*_ were 22.4, 26.8, 22.7, and 35.6 Å, respectively [29]. These DLS-measured values compare favorable to *R*_*h*_ estimated from their crystal structures [29]. A linear regression was performed on these protein standards. The *K*_*D*_ of SasG-PGR, Aap-Arpts, and SdrC-SD was inserted into the linear equation of best fit to determine the *R*_*h*_.

### R_h_ Prediction

The previously described algorithm utilizes information regarding the PPII propensity to predict *R*_*h*_ of an IDP based on the amino acid sequence [5]. To predict the hydrodynamic radius (*R*_*h*_) of a protein based on intrinsic PPII propensities, a power-law scaling relationship (Eq. 3) was used which is based on the number of residues (*N*) and the fraction of polyproline type II helix in the peptide chain (*f*_*PPII,chain*_):

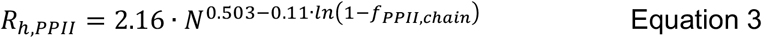

The chain propensity for PPII structure, *f*_*PPII,chain*_, is based on the experimental scale from Hilser [33] that utilized a peptide host–guest system in which the *Caenorhabditis elegans* Sem-5 SH3 domain binds a peptide in the PPII conformation. A non-interacting residue of the peptide was substituted for each amino acid before binding was measured by isothermal titration calorimetry. The value for *f*_*PPII,chain*_ in Eq. (3) was determined by Eq. (4), where *N* is the number of residues and *PPII*_*prop*_ is the PPII propensity from the Hilser scale for each amino acid in the sequence from 1 to N:

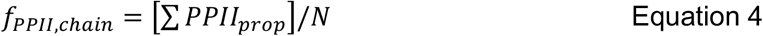

The error in the predicted *R*_*h*_ across many IDPs was found to correlate best with the net charge [29]. Based on this observation, we modified the original predictor to give Eq. (5):

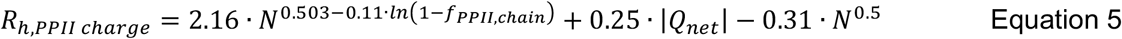

The net charge, *Q*_*net*_, is calculated from sequence as the number of lysine and arginine residues minus the number of aspartic acid and glutamic acid. Predicting *R*_*h*_ from Eq. (5) for the three proteins yields 24.4 Å for SasG-PGR, 44.1 Å for Aap-Arpts, and 22.2 Å for SdrC-SD, based on their predicted values for *f*_*PPII,chain*_ and net charge (Table S1).

### Sequence analysis of staphylococcal CWA proteins

The sequences of LPXTG-containing CWA proteins with adhesin-like function from *S. aureus* and *S. epidermidis* were submitted to the PlaToLoCo server (https://platoloco.aei.polsl.pl/) to identify potential low-complexity regions (LCRs) upstream of each protein’s LPXTG motif [80]. The server produces a consensus view from several algorithms that identify LCRs, including SEG, CAST, fLPS, and SIMPLE. We used multiple algorithms to identify a consensus region upstream of the LPXTG motif in each protein; in general, CAST provided consistent predictions for the Pro-rich proteins whereas SEG performed better at yielding consistent predictions for the SD-rich proteins. Table 5 lists the number of residues in the LCR compared to the total number of residues, which includes the sequence between the end of the LCR and the beginning of the LPXTG motif. The CIDER server (http://pappulab.wustl.edu/CIDER/) [4, 25] was used to predict Das-Pappu plot regions as well as FCR, NCPR, and omega values; *f*_*PPII*_ was calculated by the Tomasso et al. algorithm [5], SCD was calculated using the equation for SCD provided in Sawle & Ghosh [23] and the number of SD repeats, prolines, and FPR were calculated directly from the sequences.

## Abbreviations

Aap: accumulation-associated protein
CWA: cell wall-anchored
SAXS: small-angle X-ray scattering
SdrC: Serine/aspartate-rich protein C
PGR: Pro/Gly-rich region
Arpts: A-repeats
IDP: intrinsically disordered polypeptide
SasG: *S. aureus* surface protein G
SD: serine-aspartate
AUC: analytical ultracentrifugation
SEC: size-exclusion chromatography
CD: circular dichroism
PPII: polyproline type-II
SCD: sequence charge decoration
*f-*: fraction of negative residues
*f+*: fraction of positive residues
FCR: fraction of charged residues
NCPR: net charge per residue
FPR: fraction of proline residues
*f*_*PPII*_: predicted fraction of residues in the PPII conformation
kDa: kilodalton
GdnHCl: guanidinium hydrochloride
TMAO: trimethylamine *N*-oxide
TFE: trifluoroethanol
PIA: polysaccharide intercellular adhesin
PNAG: poly-*N*-acetylglucosamine
MBP: maltose-binding protein
LB: Luria-Bertani
IPTG: Isopropyl β-D-1-thiogalactopyranoside
FPLC: fast protein liquid chromatography
TEV: tobacco etch virus
LCR: low-complexity region

## Notes

**Funding:** Work was performed using funding from R01 GM094363 awarded to A.B.H. and the University of Cincinnati Graduate School Dean’s Fellowship awarded to A.E.Y. (2018-2019 AY). This research used resources of the Advanced Photon Source, a U.S. Department of Energy (DOE) Office of Science User Facility operated for the DOE Office of Science by Argonne National Laboratory under Contract No. DE-AC02-06CH11357. This project was supported by grant P30 GM138395 from the National Institute of General Medical Sciences of the National Institutes of Health. Use of the Pilatus 3 1M detector was provided by grant 1S10OD018090-01 from NIGMS. The content is solely the responsibility of the authors and does not necessarily reflect the official views of the National Institute of General Medical Sciences of the National Institutes of Health.

### Competing Interest Statement

A.B.H. serves as a Scientific Advisory Board member for Hoth Therapeutics, Inc., holds equity in Hoth Therapeutics and Chelexa BioSciences, LLC, and was a co-inventor on six patents broadly related to the subject matter of this work.

