## Supplemental Data for "Convergent behavior of extended stalk regions from staphylococcal surface proteins with widely divergent sequence patterns"

**Affiliations**:
[1] Division of Immunobiology, Cincinnati Children's Hospital Medical Center, Cincinnati, OH 45229, USA
[2] Medical Sciences Baccalaureate Program, University of Cincinnati, Cincinnati, OH 45267, USA
[3] Department of Chemistry and Biochemistry, Texas State University, San Marcos, TX 78666, USA
[4] Division of Infectious Diseases, Cincinnati Children's Hospital Medical Center, Cincinnati, OH 45229, USA
[5] Department of Pediatrics, University of Cincinnati College of Medicine, Cincinnati, OH 45229, USA

[#] Current affiliation: BioAnalysis, LLC, Philadelphia, PA 19134, USA
[†] Current affiliation: Graduate Program in Molecular Biophysics, Johns Hopkins University, Baltimore, MD 21218, USA
[‡] Current affiliation: Department of Physical Sciences, Temple College, Temple, TX 76502, USA

**Supplementary data files present in this document include:**

1. **Supplementary Tables S1—S3**

**Supplementary Tables**

**Table S1. Sequence-based parameters of IDP dataset.** The dataset is reproduced from Tomasso, et al. [1]. Parameters listed here were calculated using a program provided by Steven Whitten, based on Tomasso, et al. [1]. Shaded IDPs are from the current study. IDPs are sorted by descending $f_{PPII}$.

| IDP | N | Net charge | $R_h$ (coil) | $R_h$ (PPII) | $R_h$ (PPII charge) | $R_h$[a] (experimental) | $f_{PPII}$ |
|---|---|---|---|---|---|---|---|
| Aap-PGR | 135 | -7 | 25.64 | 38.50 | 37.84 | 37.06 | 0.5350 |
| p53(1-93) | 93 | -15 | 21.24 | 29.51 | 30.56 | 32.4 | 0.4890 |
| SasG-PGR | 69 | +7 | 18.27 | 24.56 | 24.43 | 24.8 | 0.4761 |
| p53(1-93) ALA- | 93 | -15 | 21.24 | 28.66 | 29.70 | 30.4 | 0.4581 |
| p53 TAD | 73 | -14 | 18.80 | 24.79 | 25.84 | 23.8 | 0.4500 |
| Aap-Arpts | 189 | -29 | 30.38 | 41.26 | 44.06 | 40.8 | 0.4190 |
| Securin | 202 | -1 | 31.41 | 42.57 | 40.45 | 39.7 | 0.4130 |
| PDE-γ | 87 | +4 | 20.54 | 26.51 | 25.70 | 24.8 | 0.4122 |
| Cad136 | 136 | +9 | 25.73 | 33.77 | 33.45 | 28.1 | 0.4025 |
| HIF1-α-403 | 202 | -29 | 31.41 | 42.13 | 44.86 | 44.3 | 0.4024 |
| Tau-K45 | 198 | +19 | 31.10 | 41.52 | 42.53 | 45 | 0.3988 |
| HIF1-α-530 | 170 | -10 | 28.80 | 37.81 | 37.44 | 38.3 | 0.3899 |
| Fos-AD | 168 | -16 | 28.62 | 37.17 | 37.84 | 35 | 0.3783 |
| ShB-C | 146 | -4 | 26.67 | 34.32 | 33.06 | 32.9 | 0.3764 |
| α-synuclein | 140 | -9 | 26.11 | 33.47 | 33.12 | 28.2 | 0.3744 |
| Mlph(147-403) | 260 | -28 | 35.68 | 47.00 | 49.24 | 49 | 0.3703 |
| CFTR-R-region | 189 | -5 | 30.38 | 39.18 | 37.82 | 32 | 0.3644 |
| p57-ID | 73 | -6 | 18.80 | 23.14 | 22.80 | 24 | 0.3636 |
| prothymosin-α | 110 | -43 | 23.12 | 29.02 | 34.77 | 33.7 | 0.3633 |
| LJIDP1 | 94 | +4 | 21.36 | 26.46 | 25.59 | 24.52 | 0.3565 |
| Mlph(147-240) | 97 | -15 | 21.70 | 26.85 | 27.86 | 28 | 0.3528 |
| SNAP25 | 206 | -14 | 31.73 | 40.60 | 40.70 | 39.7 | 0.3513 |
| Hdm2-ABD | 97 | -29 | 21.70 | 26.47 | 29.91 | 25.7 | 0.3345 |
| SdrC-SD | 62 | -16 | 17.31 | 20.64 | 22.15 | 21.1 | 0.3294 |
| Vmw65 | 89 | -19 | 20.78 | 25.13 | 26.90 | 28 | 0.3278 |
| p53(1-93) PRO- | 93 | -15 | 21.24 | 24.93 | 25.97 | 27.4 | 0.2832 |
| SD-30mer | 30 | -15 | 12.01 | 13.45 | 15.16 | ND[b] | 0.2700 |

[a] Reported in Å. Values in gray cells were as determined in this manuscript or [2]; values in white cells are reproduced from [1].
[b] ND, not determined.

**Table S2. The sequence of IDPs used in PPII and $R_h$ predictions.** IDP sequences (other than those from the current study - shaded) are from Tomasso, et al. supplementary material [1].

| IDP | Sequence |
|---|---|
| p53(1-93) | MEEPQSDPSVEPPLSQETFSDLWKLLPENNVLSPLPSQAMDDLMLS PDDIEQWFTEDPGPDEAPRMPEAAPPVAPAPAAPTPAAPAPAPSW PL |
| p53(1-93) ALA- | MEEPQSDPSVEPPLSQETFSDLWKLLPENNVLSPLPSQGMDDLMLS PDDIEQWFTEDPGPDEGPRMPEGGPPVGPGPGGPTPGGPGPGPS WPL |
| p53(1-93) PRO- | MEEGQSDGSVEGGLSQETFSDLWKLLGENNVLSGLGSQAMDDLML SGDDIEQWFTEDGGGDEAGRMGEAAGGVAGAGAAGTGAAGAGAG SWGL |
| p53 TAD | MEEPQSDPSVEPPLSQETFSDLWKLLPENNVLSPLPSQAMDDLMLS PDDIEQWFTEDPGPDEAPRMPEAAPRV |
| Vmw65 | GSAGHTRRLSTAPPTDVSLGDELHLDGEDVAMAHADALDDFDLDML GDGDSPGPGFTPHDSAPYGALDMADFEFEQMFTDALGIDEYGG |
| Hdm2-ABD | ERSSSSESTGTPSNPDLDAGVSEHSGDWLDQDSVSDQFSVEFEVE SLDSEDYSLSEEGQELSDEDDEVYQVTVYQAGESDTDSFEEDPEIS LADYWK |
| prothymosin-α | MSDAAVDTSSEITTKDLKEKKEVVEEAENGRDAPANGNANEENGEQ EADNEVDEEEEEGGEEEEEEEEGDGEEEDGDEDEEAESATGKRAA EDDEDDDVDTKKQKTDEDD |
| HIF1-α-403 | PAAGDTIISLDFGSNDTETDDQQLEEVPLYNDVMLPSPNEKLQNINLA MSPLPTAETPKPLRSSADPALNQEVALKLEPNPESLELSFTMPQIQD QTPSPSDGSTRQSSPEPNSPSEYCFYVDSDMVNEFKLELVEKLFAE DTEAKNPFSTQDTDLDLEMLAPYIPMDDDFQLRSFDQLSPLESSSAS PESASPQSTVTVFQ |
| Fos-AD | GSHMSVASLDLTGGLPEVATPESEEAFTLPLLNDPEPKPSVEPVKSI SSMELKTEPFDDFLFPASSRPSGSETARSVPDMDLSGSFYAADWEP LHSGSLGMGPMATELEPLCTPVVTCTPSCTAYTSSFVFTYPEADSFP SCAAAHRKGSSSNEPSSDSLSSPTLLAL |
| Mlph(147-240) | RLQGGGGSEPSLEEGNGDSEQTDEDGDLDTEARDQPLNSKKKKRL LSFRDVDFEEDSDHLVQPCSQTLGLSSVPESAHSLQSLSGEPYSED TTSLEP |
| Tau-K45 | MSSPGSPGTPGSRSRTPSLPTPPTREPKKVAVVRTPPKSPSSAKSR LQTAPVPMPDLKNVKSKIGSTENLKHQPGGGKVQIINKKLDLSNVQS KCGSKDNIKHVPGGGSVQIVYKPVDLSKVTSKCGSLGNIHHKPGGG QVEVKSEKLDFKDRVQSKIGSLDNITHVPGGGNKKIETHKLTFRENA KAKTDHGAEIVY |
| Mlph(147-403) | RLQGGGGSEPSLEEGNGDSEQTDEDGDLDTEARDQPLNSKKKKRL LSFRDVDFEEDSDHLVQPCSQTLGLSSVPESAHSLQSLSGEPYSED TTSLEPEGLEETGARALGCRPSPEVQPCSPLPSGEDAHAELDSPAA SCKSAFGTTAMPGTDDVRGKHLPSQYLADVDTSDEDSIQGPRAASQ HSKRRARTVPETQILELNKRMSAVEHLLVHLENTVLPPSAQEPTVET HPSADTEEETLRRRLEELTSNISGSSTSSE |
| p57-ID | VRTSACRSLFGPVDHEELSRELQARLAELNAEDQNRWDYDFQQDM PLRGPGRLQWTEVDSDSVPAFYRETVQV |
| PDE-γ | MNLEPPKAEIRSATRVMGGPVTPRKGPPKFKQRQTRQFKSKPPKK GVQGFGDDIPGMEGLGTDITVICPWEAFNHLELHELAQYGII |

| | |
|---|---|
| LJIDP1 | MARSFTNIKAISALVAEEFSNSLARRGYAATAQSAGRVGASMSGKMGSTKSGEEKAAAREKVSWVPDPVTGYYKPENIKEIDVAELRSAVLGKN |
| Cad136 | RLEQYTSAVVGNKAAKPAKPAASDLPVPAEGVRNIKSMWEKGNVFSSPGGTGTPNKETAGLKVGVSSRINEWLTKTPEGNKSPAPKPSDLRPGDVSGKRNLWEKQSVEKPAASSSKVTATGKKSETNGLRQFEKEP |
| α-synuclein | MDVFMKGLSKAKEGVVAAAEKTKQGVAEAAGKTKEGVLYVGSKTKEGVVHGVATVAEKTKEQVTNVGGAVVTGVTAVAQKTVEGAGSIAAATGFVKKDQLGKNEEGAPQEGILEDMPVDPDNEAYEMPSEEGYQDYEPEA |
| CFTR-R-region | GAMESAERRNSILTETLHRFSLEGDAPVSWTETKKQSFKQTGEFGEKRKNSILNPINSIRKFSIVQKTPLQMNGIEEDSDEPLERRLSLVPDSEQGEAILPRISVISTGPTLQARRRQSVLNLMTHSVNQGQNIHRKTTASTRKVSLAPQANLTELDIYSRRLSQETGLEISEEINEEDLKECLFDDME |
| SNAP25 | MAEDADMRNELEEMQRRADQLADESLESTRRMLQLVEESKDAGIRTLVMLDEQGEQLERIEEGMDQINKDMKEAEKNLTDLGKFCGLCVCPCNKLKSSDAYKKAWGNNQDGVVASQPARVVDEREQMAISGGFIRRVTNDARENEMDENLEQVSGIIGNLRHMALDMGNEIDTQNRQIDRIMEKADSNKTRIDEANQRATKMLGSG |
| ShB-C | MTLGQHMKKSSLSESSSDMMDLDDGVESTPGLTETHPGRSAVAPFLGAQQQQQQPVASSLSMSIDKQLQHPLQQLTQTQLYQQQQQQQQQQNGFKQQQQQTQQQLQQQQSHTINASAAAATSGSGSSGLTMRHNNALAVSIETDV |
| HIF1-α-530 | NEFKLELVEKLFAEDTEAKNPFSTQDTDLDLEMLAPYIPMDDDFQLRSFDQLSPLESSSASPESASPQSTVTVFQQTQIQEPTANATTTTATTDELKTVTKDRMEDIKILIASPSPTHIHKETTSATSSPYRDTQSRTASPNRAGKGVIEQTEKSHPRSPNVLSVALSQR |
| Securin | MATLIYVDKENGEPGTRVVAKDGLKLGSGPSIKALDGRSQVSTPRFGKTFDAPPALPKATRKALGTVNRATEKSVKTKGPLKQKQPSFSAKKMTEKTVKAKSSVPASDDAYPEIEKFFPFNPLDFESFDLPEEHQIAHLPLSGVPLMILDEERELEKLFQLGPPSPVKMPSPPWESNLLQSPSSILSTLDVELPPVCCDIDI |
| Aap-PGR | AEPGKPAEPGKPAEPGKPAEPGTPAEPGKPAEPGTPAEPGKPAEPGKPAEPGKPAEPGKPAEPGTPAEPGTPAEPGKPAEPGTPAEPGKPAEPGTPAEPGKPAESGKPVEPGTPAQSGAPEQPNRSMHSTDNKNQ |
| SasG-PGR | PKDPKGPENPEKPSRPTHPSGPVNPNNPGLSKDRAKPNGPVHSMDKNDKVKKSKIAKESVANQEKKRAE |
| Aap-Arpts | NNEAPQMSSTLQAEEGSNAEAPQSEPTKAEEGGNAEAAQSEPTKAEEGGNAEAPQSEPTKAEEGGNAEAAQSEPTKTEEGSNVKAAQSEPTKAEEGSNAEAPQSEPTKTEEGSNAKAAQSEPTKAEEGGNAEAAQSEPTKTEEGSNAEAPQSEPTKAEEGGNAEAPQSEPTKTEEGGNAEAPNVPTIKA |
| SdrC-SD | SDSDSDSDSDSDSDSDSDSDSDSDSDSDNDSDSDSDSDSDAGKHTPAKPMSTVKDQHKTAKA |
| SD-30mer | SDSDSDSDSDSDSDSDSDSDSDSDSDSDSD |

**Table S3: The sequence of low-complexity regions from Staphylococcal CWA proteins from Table 5.** Sequences start at the beginning of the consensus LCR region identified by the PlaToLoCo server [3] and extend through the sequence immediately upstream of the LPXTG sortase motif. See Materials and Methods for further details.

| Protein | Sequence |
|---|---|
| *SD-rich LCRs* | |
| SdrC | TSDSDSDSDSDSDSDSDSDSDSDSDSDSDSDSDSDSDSDSDSDSDSDSDSNSD<br>SDSDSDSDSDSDSDSDSDSDSDSDSDSDSDSDSDSDSDSDSDSDSDSDSDSDS<br>DSDSDSDSDSDSDSDSDSDSDSDSDSDSDSDSDSDSDSDSDSDSDSDSDSDSD<br>NDSDSDSDSDSDAGKHTPAKPMSTVKDQHKTAKA |
| SdrD | TSDSDSDSDSDSDSDSDSDSDSDSDSDSDSDSDSDSDSDSDSDSDSDSDSDSDSD<br>SDSDSDSDSDSDSDSDSDSDSDSDSDSDSDSDSDSDSDSDSDSDSDSDSDSDS<br>DSDSDSDSDSDSDSDSDSDSDSDSDSDSDSDSDSDSDAGKHTPVKPMSATKDH<br>HNKAKA |
| SdrE | TSDSDSDSDSDSDSDSDSDSDSDSDSDSDSDSDSDSDSDSDSDSDSDSDSDSDSD<br>SDSDSDSDSDSDSDSDSDSDSDSDSDSDSDSDSDSDSDSDSDSDSDSDSDSDS<br>DSDSDSDSDSDSDSDSDSDSDSDSDSDSDSDSDSDSDSDSDSDSDSDSDSDSD<br>AGKHTPVKPMSTTKDHHNKAKA |
| SdrF (*S. epi*) | TSDSDSDSDSDSDSDSDSDSDSDSDSDSDSDSDSDSDSDSDSDSDSDSDSDSDSD<br>SDSDSDSDSDSDSDSDSDSDSDSDSDSDSDSDSDSDSDSDSDSDSDSDSDSDS<br>DSDSDSDSDSDSDSDSDSDSDSDSDSDSDSDSDSDSDSDSDSDSDSDSDSDSD<br>NDSDSDSDSDSDAGKHTPAKPMSTVKDQHKTAKA |
| SdrG (*S. epi*) | TSDSDSDSDSDSDSDSDSDSDSDSDSDSDSDSDSDSDSDSDSDSDSDSDSDSDSD<br>SDSDSDSDSDSDSDSDSDSDSDSDSDSDSDSDSDSDSDSDSDSDSDSDSDSDS<br>DSDSDSDSDSDSDSDSDSDSDSDNDSDSDSDSDSDAGKHTPAKPMSTVKDQHK<br>TAKA |
| Pls | DSDADSDSDADSDSDADSDSDADSDSDADSDSDSDSDSDSDSDADSDSDSD<br>SDSDADSDSDADSDSDADSDSDSDADSDSDSDSDADSDSDADSDSDADS<br>DSDSDSDSDADSDSDSDSDSDADSDSDADSDSDADSDSDADSDSDSDSDSDAD<br>SDSDADSDSDADSDSDADSDSDSDSDSDADSDSDSDSDSDADSDSDADSDS<br>DSDADSDSDADSDSDADSDSDADSDSDSDSDSDADSDSDADSDSDADSDSDAD<br>SDSDSDSDSDSDSDSDADSDSDSDSDSDADRDHNDKTDKPNNKE |
| ClfA | VPEQPDEPGEIEPIPEDSDSDPGSDSGSDSNSDSGSDSGSDSTSDSGSDSASDS<br>DSASDSDSASDSDSASDSDSASDSDSDNDSDSDSDSDSDSDSDSDSDSDSDSD<br>SDSDSDSDSDSDSDSDSDSDSDSDSDSDSDSDSDSDSDSDSDSDSDSDSDSDS<br>DSDSDSDSDSDSDSDSDSDSDSDSDSDSDSDSDSDSDSDSDSDSDSDSDSDSD<br>SDSDSDSDSDSASDSDSDSDSDSDSDSDSDSDSDSDSDSDSDSDSDSDSDSES<br>DSDSESDSDSDSDSDSDSDSDSDSDSASDSDSGSDSDSSSDSDSESDSNSD<br>SESGSNNNVVPPNSPKNGTNASNKNEAKDSKEP |
| ClfB | VDPEPSPDPEPEPTPDPEPSPDPEPEPSPDPDPDSDSDSDSGSDSDSGSDSDSE<br>SDSDSDSDSDSDSDSDSESDSDSESDSDSDSDSDSDSDSDSESDSDSDSDSDS<br>DSDSDSESDSDSESDSESDSDSDSDSDSDSDSDSDSDSDSDSDSDSDSDSDSD<br>SESDSDSDSDSDSDSDSDSDSDSDSDSDSDSDSDSDSDSDSDSDSDSDSDSDS<br>DSDSDSDSDSDSDSDSDSDSDSDSDSDSDSDSDSDSDSDSRVTPPNNEQKAPS<br>NPKGEVNHSNKVSKQHKTDA |
| SesJ (*S. epi*) | FEDSESDSSSESESDSESHSDSESHSDSESTSESDSESHSDSESTSESDSESHS<br>DSESDSDSESTSESDSESHSDSESDSDSESTSESDSESHSDSESHSDSESTSES<br>DSESHSDSESDSDSESTSESDSESHSDSESHSDSESTSESDSESHSDSESDSDS<br>ESTSESDSESHSDSESDSDSESTSESDSESHSDSESDSDSESTSESGSESHSNS<br>E |

| | |
|---|---|
| *Pro-rich LCRs* | |
| Aap [a] (*S. epi*) | PTKAEPGKPAEPGKPAEPGKPAEPGTPAEPGKPAEPGTPAEPGKPAEPGKPAEP<br>GKPAEPGKPAEPGTPAEPGTPAEPGKPAEPGTPAEPGKPAEPGTPAEPGKPAES<br>GKPVEPGTPAQSGAPEQPNRSMHSTDNKNQ |
| SasG | PKDPKGPENPEKPSRPTHPSGPVNPNNPGLSKDRAKPNGPVHSMDKNDKVKKS<br>KIAKESVANQEKKRAE |
| CNA | PEKPNKPIYPEKPKDKTPPNKPDHSNKVRPTPPDEPSKVDKVDQPKDNKTKPENP<br>LKE |
| FnbpA | PPIVPPTPPTPEVPSEPETPTPPTPEVPSEPETPTPPTPEVPSEPETPTPPTPEVPA<br>EPGKPVPPAKEEPKKPSKPVEQGKVVTPVIEINEKVKAVAPTKKPQSKKSE |
| FnbpB | PPIVPPTPPTPEVPSEPETPTPPTPEVPSEPETPTPPTPEVPTEPGKPIPPAKEEPK<br>KPSKPVEQGKVVTPVIEINEKVKAVVPTKKAQSKKSE |
| *Other LCRs* | |
| SraP (SasA) | MSGSQSISDSTSTSMSGSTSTSESNSMHPSDSMSMHHTHSTSTSRLSSEATTST<br>SESQSTLSATSEVTKHNGTPAQSEKR |
| FmtB (SasB) | NNKATQNDGANASPATVSNGSNSANQDMLNVTNTDDHQAKTKSAQQGKVNKAK<br>QQAKT |
| SasC | DTAIGQIDQDRSNAQVDKTASLNLQTIHDLDVHPIKKPDAEKTINDDLARVTALVQN<br>YRKVSDRNKADALKAITALKLQMDEELKTARTNADVDAVLKRFNVALSDIEAVITEK<br>ENSLLRIDNIAQQTYAKFKAIATPEQLAKVKVLIDQYVADGNRMIDEDATLNDIKQH<br>TQFIVDEILAIKLPAEATKVSPKEIQPAPKVCTPIKKEETHESRKVEKE |

[a] The Aap sequence listed here is based on the consensus identification of the LCR region by the PlaToLoCo server [3], as for all other sequences in Table 5. This sequence differs slightly from the Aap construct used for experimental approaches (compare to Figure 1).
